# DuReS: An R package for denoising experimental tandem mass spectrometry-based metabolomics data

**DOI:** 10.1101/2024.09.16.613198

**Authors:** Shayantan Banerjee, Prajval Nakrani, Aviral Singh, Pramod P. Wangikar

## Abstract

Mass spectrometry-based untargeted metabolomics is a powerful technique for profiling small molecules in biological samples, yet accurate metabolite identification remains challenging. One of the primary obstacles in processing tandem mass spectrometry data is the prevalence of random noise peaks, which can result in false annotations and necessitate labor-intensive verification. A common method for removing noise from MS/MS spectra is intensity thresholding, where low-intensity peaks are discarded based on a user-defined cutoff or by analyzing the top “N” most intense peaks. However, determining an optimal threshold is often dataset-specific and may retain many noisy peaks. In this study, we hypothesize that true signal peaks consistently recur across replicate MS/MS spectra generated from the same precursor ion, unlike random noise. An optimal recurrence frequency of 0.12 (95% CI: 0.087-0.15) was derived using an open-source metabolomics dataset, which enhanced the dot product score between the experimental and library spectra by 66% post-denoising and resulted in a median signal and noise reduction of 5.83% and 99.07%, respectively. Validated across multiple metabolomics datasets, our denoising workflow significantly improved spectral matching metrics, leading to more accurate annotations and fewer false positives. Available freely as an R package, Denoising Using Replicate Spectra (DuReS) (https://github.com/BiosystemEngineeringLab-IITB/dures) is designed to remove noise while retaining diagnostically significant peaks efficiently. It accepts mzML files and feature lists from standard global untargeted metabolomics analysis software as input, enabling users to seamlessly integrate the denoising pipeline into their workflow without additional data manipulation.

## Introduction

Mass spectrometry-based untargeted metabolomics provides a high-throughput approach for quantifying small molecules in complex mixtures. Developing reproducible preprocessing strategies is crucial for reliably identifying true metabolic features in these experiments. A key challenge in mass spectra preprocessing is reducing spectral complexity while effectively identifying ions in untargeted analyses. Several methods have been developed to denoise spectra from LC-MS experiments based on certain characteristics that differentiate between signal and noisy peaks [4-7]. Although these techniques have significantly improved peak detection from LC-MS data, few have concentrated on reducing complexity in tandem mass spectrometry (MS/MS) data, which can offer improved specificity and sensitivity over LC-MS by utilizing two stages of mass analysis, allowing for more accurate identification of compounds. In a typical MS/MS spectrum, only a small fraction of peaks are critical for metabolite annotation, with the majority arising from background noise and contaminants. Since metabolite identification relies on the similarity between experimental and reference spectra, these biologically irrelevant peaks or noise ions lower the signal-to-noise ratio and increase the likelihood of false positive annotations [1-3]. To address the challenges caused by noise, a preprocessing step is typically applied to denoise the MS/MS spectra and detect signal peaks, ensuring more accurate metabolite identification.

Random noise originating from instruments is characterized by small peaks uniformly distributed throughout the spectrum [8]. One of the most commonly used techniques to remove random noise from MS/MS spectra is intensity thresholding, wherein peaks with intensities less than a user-defined threshold are discarded from the spectrum. Sometimes, a fixed percentage of the maximum peak intensity and/or limiting the analysis to the top “N” most intense peaks is employed to reduce spectrum complexity [9,10]. However, an optimal threshold value is often dataset-specific and hard to determine [11]. Automated peak elimination techniques [12] address this issue by iteratively discarding low-intensity noise peaks based on user-defined thresholds for noise fragment ratio, relative standard deviation (RSD), and peak intensity. Another preprocessing technique, binning, reduces noise peaks in the spectrum by grouping adjacent peaks within a given m/z window. However, the window width influences the quality of denoising, and a large value may accidentally remove signal peaks [13].

Multiple studies have shown that acquiring replicate MS/MS spectra from the same precursor ion improves the signal-to-noise ratio and enhances confidence in metabolite annotation [14-16]. Averaging replicate spectra to generate consensus spectra is vital for creating high-quality spectral libraries, such as NIST, by filtering out spurious peaks [17]. However, these studies mainly focus on curating spectral libraries and offer limited open-source tools for denoising experimental tandem mass spectra. We hypothesized that signal peaks, even those corresponding to low-intensity fragments, recur frequently. In contrast, noisy peaks, regardless of intensity, do not recur. Averaging replicate spectra from curated metabolic features could help establish an optimal frequency threshold for identifying true signals. Therefore, a voting scheme can retain diagnostically significant low-intensity fragments that appear consistently, otherwise lost in intensity thresholding or top-N techniques. In this study, we introduce the Denoising using Replicate Spectra (DuReS) R package, which leverages open-source metabolomics datasets to derive and validate an optimal frequency threshold for denoising experimental MS/MS data and provides a detailed protocol for this preprocessing step.

## Experimental Section

### Proposed algorithm

The methodology consists of two modules: (i) “Tuning module,” which determines an optimal frequency threshold to classify peaks in an MS2 spectrum as either signal or noise. Following denoising, the spectrum is matched against a reference library, with accuracy and noise reduction evaluated using various metrics (Figure S3); and (ii) “Testing module,” which validates the score cutoff on unannotated features by comparing pre-and post-denoising annotations, flagging low-confidence hits as false positives, and utilizing mirror plots to compare experimental and reference spectra visually. The problem is formally defined in Section 3 of Supporting Information.

### Tuning Module

#### Extraction of MS/MS Spectra and preparation of an annotated list of metabolites before denoising

The tuning module starts with a directory containing mzML files and a list of MS/MS annotated features characterized by its ID, precursor m/z value, start and end retention time (RT) in minutes, the spectrum with best pre-denoising matching scores (see section 1 of supporting information), and the corresponding reference database entry (Spec_best_, Ref_best_) and annotation. The feature lists, annotations, and spectra-reference pair (Spec_best_, Ref_best_) were generated using our in-house web-based software suite, MSOne (https://msone.claritybiosystems.com/). Only metabolites with matching scores greater than or equal to 700 were included in the tuning module. Duplicate and unannotated features were not considered for further analysis. Alternatively, the user can use any peak picking and alignment software to generate the input feature list for our tool. To extract MS/MS spectra from mzML files, we introduced a fixed mass tolerance of 5 ppm. The corresponding tolerance in daltons was calculated using the formula below (5).

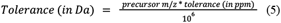

Given the RT and m/z ranges, two functions from the Spectra package in R, `filterRt()` and `filterPrecursorMzRange()`, were used to extract the MS2 spectra for every feature in the list.

#### Generating aggregate spectra and labeling peaks with consistency scores

For a given feature, individual spectra are sorted in decreasing order based on the Total Ion Content (TIC) values, and the top 80% are combined to form an inter-spectra aggregate. The consistency factor (*Cf*) for every representative peak (Section 3 of supporting information) belonging to the aggregate, its mean m/z, and intensity is reported, and the same is used to label the peaks from the individual spectrum (Spec_best_).

#### Deriving an optimal consistency factor cutoff

Denoised spectra for each feature were generated using consistency factor thresholds from 0 to 1 in 0.01 increments, incorporating peaks that met the criterion. These spectra were matched to the reference entry (Supporting information, Section 2), resulting in multiple matches evaluated using metrics (Supporting Information, Section 1) before and after denoising. To identify the optimal consistency factor (*Cf*\*), a multiobjective optimization was performed to balance Signal Reduction (SR) and Noise Reduction (NR) using a Pareto front, where each point represents a trade-off between the two objectives. *Pareto Front*= {(*SR _i_, NR_i_*)| *i* = 1, 2,… *n*} where *n* is the total number of points on the Pareto front. A single solution from the Pareto front is selected as follows. Let’s denote the set of Pareto optimal solutions as *P*_*i*_, and each solution in *P*_*i*_ is represented as *p*_*i*_. The matching score of each solution *p*_*i*_ is denoted as *MS(p*_*i*_). We define *p\** with the maximum similarity score among all the Pareto optimal solutions.

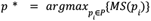

Among all Pareto efficient solutions, representing the best tradeoff in percentage signal and noise reduction, the optimal consistency factor (*Cf*\*) was selected with the best matching score for that feature. The `psel` function from the `rPref` package in R was used to determine the optimal consistency score. The entire workflow for the tuning module is outlined in Figure S1.

### Testing Module

#### Consensus building and library matching

During the tuning phase, we identified the optimal frequency cutoff by analyzing high-quality MS2 features, focusing on annotated metabolites with matching scores above 700. The optimal cutoff was then applied to the complete set of unannotated MS2 features in the testing phase. We retained features with at least ten spectra for analysis in the testing phase. To improve computational efficiency, spectra were ranked by Total Ion Chromatogram (TIC) values, and the top 80% were retained. After excluding fragments with zero intensities, an inter-spectra aggregate (Supporting information; Section 3) was created, and consistency scores for representative peaks were calculated as described previously. For every feature, the top 50 spectra from the top 80% list were selected, and their corresponding fragments were labeled with the consistency factors calculated using the aggregate.

Next, an intra-spectra peak grouping was performed for each reference spectrum. The 50 experimental spectra corresponding to a particular MS2 feature were compared against the processed reference library. A consistent mass-to-charge (m/z) tolerance of 0.05 Da was applied during the matching process. Subsequently, the matches were organized according to their matching scores, excluding those scoring below 300. Annotations are evaluated based on their prevalence across the 50 scans, and the annotations with the highest occurrence frequency are attributed to the respective feature. Only annotations exhibiting an occurrence frequency of at least 30% were considered valid. Similarly, features that exhibited forward reverse and modified dot products below 0.25 and less than two fragments matching with the reference were removed from the analysis.

#### Applying a fixed optimal frequency cutoff for removing noisy fragments

Fragments from the 50 experimental spectra with consistency factors meeting the optimal threshold criteria are retained for every unannotated feature. We set lower and upper cutoffs for signal presence count at 3 and 50, respectively, to derive denoised spectra. For instance, if an unannotated MS2 feature has 1000 spectra and a consistency factor cutoff of 0.12, the resulting signal presence count is 120 (1000*0.12). However, the probability of observing noisy fragments recurring across all 120 scans is extremely low. Therefore, we clip the cutoff for signal presence count to 50. Similarly, the signal presence count for an MS2 feature with only ten spectra is 1.2 (10*0.12). However, to ensure compatibility with our algorithm, we clip this to 3. The denoised spectra obtained using the above cutoffs are then compared against the combined reference library (Supporting Information, Section 2), and several matching metrics (Supporting information, Section 1, Figure S3) are derived. An approach similar to the “consensus building and library matching” section from the “Testing Module” was adopted to derive annotations from the matches obtained using the 50 denoised spectra. The entire workflow for the testing module is outlined in Figure S2.

## Results and Discussion

The details of all the datasets used in this study are listed in Table 1. To determine the optimal consistency factor, we used the World Trade Centre - Lung Injury dataset [18] (Metabolomics Workbench ID: ST001711), which includes LC-MS/MS data of 248 individuals exposed to particulate matter. The tuning dataset, consisting of 42 high-quality MS2 features with raw matching scores above 700, was used to derive the optimal consistency factor. Validation was performed using four datasets, including the remaining unannotated features not included in the tuning phase from the WTC-LI dataset, data from plasma and swab samples (MTBLS2291, MTBLS2349) [19] derived from COVID-19 patients from India, a study on plasma, and serum eicosadomics (ST003050) [20], and a dataset used to perform a benchmarking analysis [21] to evaluate different software packages for untargeted metabolomics analysis (XCMS dataset ID: 1197236).

**Table 1:**
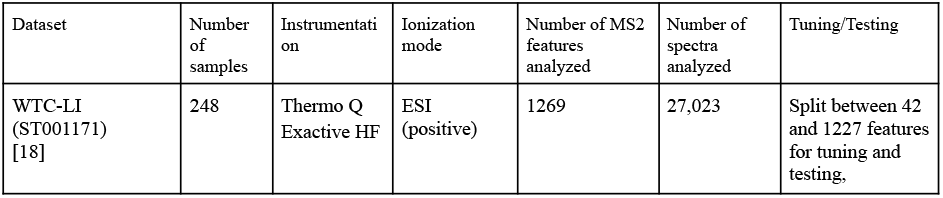
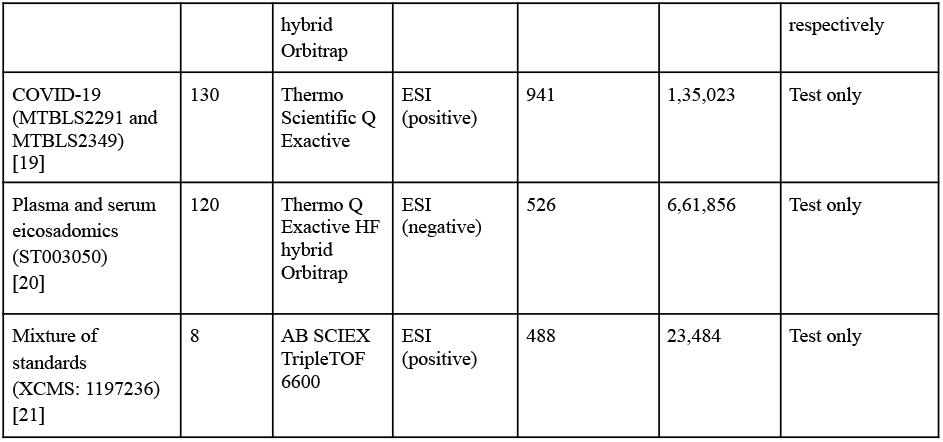
Details of the datasets used in this study.

### Enhanced Matching Scores During Tuning Phases Post-Denoising

Using 42 metabolites that met our inclusion criteria, we performed spectral matching before and after denoising with the combined reference library (Table 2; Figure 1A-H). The median number of spectra per feature was 381.5 (95% CI: 266.16-496.84). After denoising, we observed a significant increase in the median matching scores, the fraction of matching fragments, and forward and modified dot products (Wilcoxon test; p < 0.05). However, no significant differences were found for reverse dot products or the number of matching fragments.

**Table 2:**
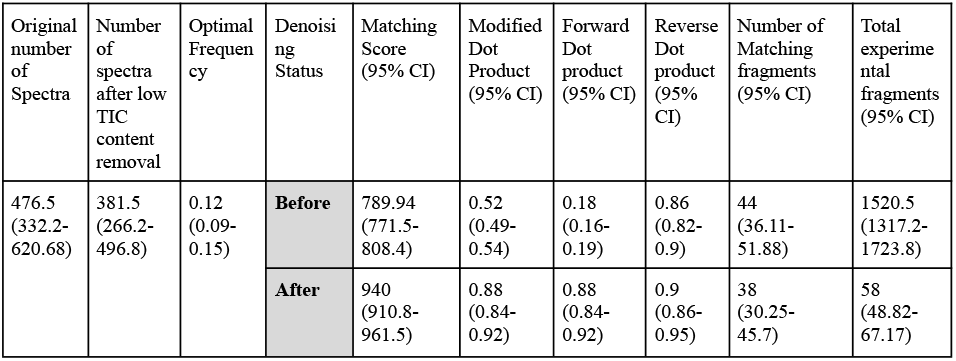
Median values of spectral matching metrics during the tuning phase (42 MS2 features, matching score before denoising >= 700)

**Figure 1A-H:.**
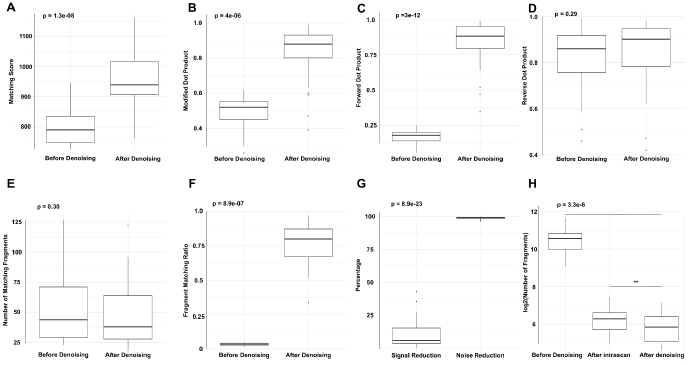
Matching metrics before and after denoising during the tuning phase.

The tuning phase achieved median signal and noise reductions of 5.83% (95% CI: 2.75-8.9%) and 99.07% (95% CI: 98.87-99.08%), respectively. The median logarithms (base 2) of fragments per spectrum per feature were 10.57 (95% CI: 10.37-10.77) before denoising, 6.3 (95% CI: 6.1-6.5) after intra-spectrum peak grouping, and 4.06 (95% CI: 3.9-4.20) post optimal consistency factor cutoff. Significant differences (Wilcoxon test; p<0.05) were observed across all three stages of denoising. The median matching score increased by 21.43% (95% CI: 19.47-23.40%), with a modified dot product of 66.22% (95% CI: 59.03-73.41%). Using multi-objective optimization, we derived Pareto fronts for all 42 features, pinpointing the optimal frequency cutoff that achieves the best trade-off between minimizing signal loss and maximizing noise reduction (Figure S5). The median optimal consistency factor cutoff was 0.12 (95% CI: 0.087-0.147).

### Enhanced Matching Scores During Testing Phases Post-Denoising

The median number of spectra per feature was 368 (95% CI: 289.6-446.4) for the WTC-LI dataset, 54 (95% CI: 46.47-61.52) for the COVID-19 dataset, 201 (95% CI: 65-336) for the plasma and serum eicosadomics dataset, and 48 (95% CI: 45.76-50.24) for the mixture of standards dataset. Applying a fixed frequency cutoff improved metrics for matching quality across all datasets (Supporting Information: Table S1A; Figure S4(i-iv)). Significant increases in median matching scores were observed for the WTC-LI and mixture of standards datasets (Wilcox test; p<0.05), with no significant changes in the other datasets. Fragment matching ratio, forward, and modified dot products increased significantly across all datasets (Wilcox test; p<0.05). However, the median number of experimental fragments decreased in all datasets, with significant reductions only in the WTC-LI and COVID-19 datasets. A significant decrease in the number of experimental fragments was observed across all three stages of denoising (before, after intra-spectrum, and after frequency thresholding) (Table S1B)

### Fixed Frequency Cutoff-Based Denoising Yields Superior Matches Compared to Fixed Intensity Cutoff

We compared our fragment frequency-based denoising technique against a conventional method that removes fragments below a fixed intensity threshold (x = 0.1, 0.5, 1, 5, and 10 percent of the spectrum’s maximum intensity). Our approach consistently outperformed the standard method in matching scores and dot products (modified, forward, and reverse) while retaining more matching fragments after denoising (Figure S4 (i-iv)).

### Use cases: The annotation remained unchanged, but the matching score improved after denoising

After denoising, several metabolites retained the same annotation for all four validation sets, but their match quality significantly improved. This enhancement is evident in the substantial increase in matching scores and modified dot products. For instance, after applying our denoising algorithm on L-γ-Glutamyl-L-glutamic Acid (precursor m/z: 277.1 Da) and Harmol (precursor m/z: 199.1 Da), the matching scores, forward, reverse, and modified dot products increased from 621.897, 0.06, 0.465, and 0.262 to 883.956, 0.975, 0.862, 0.918, and from 712.81, 0.05, 0.75, 0.4 to 983.06, 0.895, 0.908, and 0.902, respectively (Figure 2A-B, E-F). Similar observations were made for the metabolites cytosine, lauroyl-carnitine, and cholic acid (Figure S6-S8).

**Figure 2 A-B, E-F:**
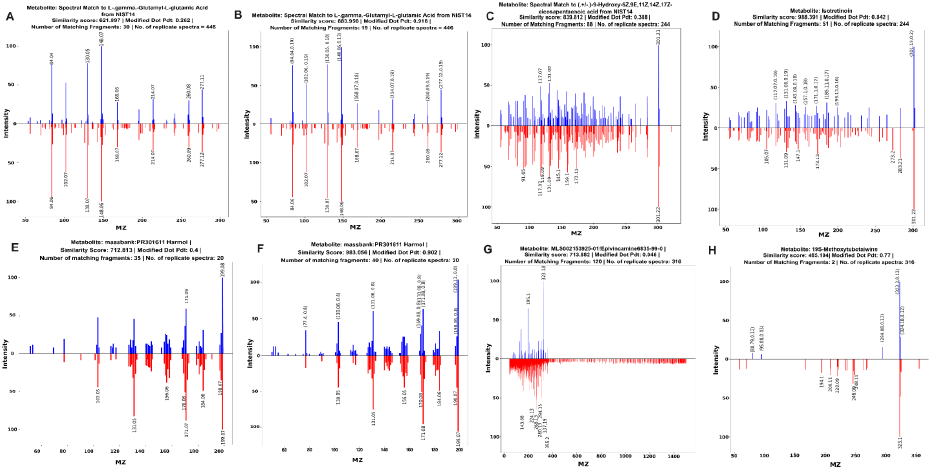
Before (Fig 2A, 2E) and after (Fig 2B, 2F) denoising mirror plots for L-.gamma.-Glutamyl-L-glutamic Acid and Harmol displaying the experimental and library spectrum match. Annotation remained the same after denoising, but the match quality improved. **Figure 2C-D:** Before (Fig 2C) and after (Fig 2D) denoising mirror plot for Isotretinoin, where the annotation changed and the match quality improved after denoising. **Figure 2G-H:** False positive annotation where the matching score dropped drastically after denoising.

### The annotation changed, and the matching score improved after denoising

Figure 2C-D shows the scenario where, after denoising, the annotation of the metabolite changed. Both the reference and the experimental spectrum changed after denoising, and the matching score, forward, reverse, and modified dot products increased from 839.812, 0.125, 0.65, and 0.388 to 988.391, 0.875, 0.81, and 0.842, respectively. The annotation changed from (.+/-.)-9-Hydroxy-5Z,9E,11Z,14Z,17Z-eicosapentaenoic acid to Isotretinoin. Similarly, we observe the annotation for the precursor ion (m/z: 277.22 Da) change from 9S-Hydroxy-10E,12Z,15Z-octadecatrienoic acid to 13-Keto-9Z,11E-octadecadienoic acid (Figure S9)

### The annotation changed, and the matching score decreased after denoising (probable false positives)

The presence of numerous noisy fragments led to inflated matching scores as some fragments aligned randomly between the experimental and the noisy reference spectrum. After denoising, these inflated scores dropped significantly. For instance, the matching score for the precursor ion (m/z: 355.209) decreased from 713.88 to 448.104, leaving only two matching fragments (Figure 2E-F). A similar trend was observed for other features with m/z values (Figure S10), which were subsequently flagged as false positive annotations.

### General recommendations for identifying false positive annotations using our algorithm

A matching score-wise metrics breakdown revealed that features with after-denoising matching scores above 800 consistently lead to reliable annotations (Tables S2-S5). For features with matching scores between 600 and 800, further visual inspection using mirror plots is necessary, along with evaluating metrics like the number of matching fragments and dot products. A modified dot product of *≥*0.8 with *≥*10 matching fragments typically suggests a strong match. Conversely, features with scores below 500 can be flagged as likely false positives.

## Software implementation

DuReS is an open-source R package (v4.0+) available under the Apache 2.0 license. It can be installed directly from GitHub (https://github.com/banerjeeshayantan/dures) using the ‘install_github()’ function from the ‘devtools’ package [22]. The tool processes a folder ofmzML files and a feature list obtained from any software for untargeted metabolomics preprocessing, consisting of m/z and RT values, outputting denoised spectra for those features. Comprehensive documentation and test cases are included within the package.

## Conclusion

We present an open-source R package to denoise experimental tandem mass spectrometry-based metabolomics data. Instead of relying on the conventional intensity threshold method for removing noisy fragments, our approach calculates the occurrence frequency of fragments across replicate MS/MS spectra to identify and label noise. We derived an optimal frequency threshold using metabolites with high-quality annotations from an open-source untargeted metabolomics dataset. We determined that a consistency factor cutoff of 0.12 (95% CI: 0.087-0.147) effectively removes noise from experimental tandem mass spectra, as validated across four independent datasets. Our results significantly improve matching quality metrics compared to traditional intensity-based denoising methods. Our validation experiments suggest that the optimal frequency threshold reliably retains most signal peaks, enhancing matching scores and reducing potential false positives across various experimental metabolomics datasets. However, further validation with more diverse datasets from different instrument types and experimental designs is necessary to confirm the stability and robustness of our approach.

## Notes

### Competing Interest Statement

The authors have declared no competing interest.

