## Supplementary Information for "DuReS: An R package for denoising experimental tandem mass spectrometry-based metabolomics data"

Shayantan Banerjee[1], Prajval Nakrani[2], Pramod P. Wangikar*[1,2,3]
[1]Centre for Digital Health, Indian Institute of Technology Bombay, Mumbai, India
[2]Clarity Bio Systems India Pvt. Ltd., Mumbai, India
[3]Department of Chemical Engineering, Indian Institute of Technology Bombay, Mumbai, India

**Table of Contents**

**Section 1. Metrics to judge spectral matches------------------------------------------------------------ 1**
**Section 2: Reference Library Preparation------------------------------------------------------------------ 2**
**Section 3: Problem formulation--------------------------------------------------------------------------- 3**
**Table S1A: Median values of spectral matching metrics during the validation phase------- 5**
**Table S1B: Number of fragments during three stages of denoising in the validation phase 6**
**Table S2 - Median values of matching metrics across different matching score ranges-(WTC-LI)------------------------------------------------------------------------------------------------- 7**
**Table S3 - Median values of matching metrics across different matching score ranges-(COVID-19)-------------------------------------------------------------------------------------------------8**
**Table S4 - Median values of matching metrics across different matching score ranges-(ST003050)-------------------------------------------------------------------------------------------------9**
**Table S5 - Median values of matching metrics across different matching score ranges-(benchmarking)-------------------------------------------------------------------------------------------- 10**
**Figure S1 - Tuning phase workflow to derive optimal frequency threshold—---------------- 11**
**Figure S2 - Testing phase workflow to evaluate the optimal frequency threshold —------ 12**
**Figure S3 - Graphical representation of formulation of matching metrics used in this study —-----------------------------------------------------------------------------------------------------13**
**Figure S4 (i-iv)- Comparison of matching metrics between different denoising approaches using the WTC-LI, COVID-19, ST003050, Benchmarking datasets —------ 14-17**
**Figure S5 - Plots showing the tradeoff between signal and noise reduction in the tuning phase —---------------------------------------------------------------------------------------------------18**
**Figure S6-S10 - Case studies —------------------------------------------------------------------- 19-23**

#### Section 1. Metrics to judge spectral matches

An essential aspect of annotating unknown MS2 spectra involves the computation of similarity scores between experimental and library spectra. In this study, we used the following metrics to judge the quality of spectral matches. A pictorial representation of the metrics is available in Figure S3.

- Number of matching fragments: Number of matching peaks between the reference and the experimental spectrum within a given m/z tolerance (0.05 Da).

- Forward dot product: each spectrum is treated as an ordered list of peak intensities, and the cosine of the angle between the experimental and the reference spectra is reported. Fragment ions of the experimental spectra define the vector space for the dot product. In the case of a noisy experimental spectrum, the forward dot product is usually small.

- Reverse dot product: uses the same formula as forward dot product except that the fragment ions of the reference spectra define the vector space for dot product. In the case of a noisy experimental spectrum, the reverse dot product can be high, leading to false positive annotations.

- Modified dot product: is the average between forward and reverse dot products.

- Matching Score: we developed a new scoring system that considers the number of matching fragments and the modified dot product to evaluate the overlap between two spectra. It is given by,

$$Matching\ Score\ =\ 5\ *\ [Mod.\ Dot\ Pdt.\ \ *\ 100\ +\ 20\ *\ log_2(max(NMF, 1))]$$

where NMF = number of matching fragments

- $Percent\ signal\ reduction\ =\ \frac{Q-P}{Q}\ *\ 100$
- $Percent\ noise\ reduction\ =\ \frac{[(M-Q)-(N-P)]}{M-Q}\ *\ 100$

where,
Q = number of peaks matching between experimental and reference spectrum before denoising;
P = number of peaks matching between experimental spectra matching with reference after denoising;
M = number of peaks in the experimental spectrum before denoising;
N = number of peaks in the experimental spectrum after denoising

- Fragment Matching Ratio: Fraction of fragments matching with the reference. Denotes the noise content in the dataset. A high fragment-matching ratio corresponds to low noise content.

$$Fragment\ Matching\ ratio\ =\ \frac{Number\ of\ matching\ fragments}{Total\ number\ of\ fragments\ in\ the\ spectrum}$$

### Section 2: Reference Library Preparation

A combined reference library with MS/MS data from both ionization modes was curated by consolidating entries from MSDial, HMDB, GNPS, and Lipid Blast databases. Each entry was assigned a unique SPLASH key using the splashR package, with duplicates removed and intra-spectrum peak grouping performed. The m/z and intensity values for each metabolite were extracted and stored, with a fixed m/z tolerance of 0.05 Da applied for matching MS/MS spectra. The best matches between the experimental and library spectra were visualized using mirror plots for both the tuning and the testing modules. Peak intensities were square root transformed to avoid a strong dependence on high-intensity peaks.

### Section 3: Problem formulation

For a given MS1 feature $F$, let there be $N$ replicate MS2 spectra, $S = \{s_1, s_2, ... s_N\}$. Each spectrum $s_i$ can be represented as a series of peaks, $s_i = \{p_{i1}, p_{i2}, ... p_{in}\}$, where $n = |s_i|$. The j[th] peak of the i[th] spectrum $p_{ij}$ can be described as a pair $(m/z_{ij}, intensity_{ij})$. We define "consistency factor" as the fraction of times a given peak $p_{ij}$ recurs across $N$ replicate MS2 spectra within a given m/z tolerance. Formally, it is represented as follows,

Given:

- N spectra $S = \{s_1, s_2, ... s_N\}$
- Peak $p_{ij}$ with mass value $m/z_{ij}$ and intensity $I_{ij}$
- The m/z tolerance $t$ for considering two peaks as a "match."

**Intra-spectrum peak grouping:** Within a given spectrum, peaks with non-zero intensities whose m/z lie within a predefined tolerance value of one another are merged using the formula shown below:

For peaks $p_{ij}$ $and$ $p_{ik}$ belonging to the same spectrum $i$, if $|m/z_{ij} - m/z_{ik}| \leq t$, then

$$m/z_{merged} = \frac{m/z_{ij} + m/z_{ik}}{2} \quad (1)$$

$$I_{merged} = I_{ij} + I_{ik} \quad (2)$$

**Inter-spectra aggregate (creating representative spectrum):** All intra-spectrum grouped spectra $S' = \{s'_1, s'_2, ... s'_N\}$ from the previous step are combined to represent a single representative spectrum known as the aggregate. Across spectra, peaks were merged if they lay within a tolerance $t$ of one another, and the aggregate spectrum was created using the average mass and intensities as shown below:

Given:

- N intra-spectrum grouped spectra $S' = \{s'_1, s'_2, ... s'_N\}$

- Each grouped spectrum $s'_i = \{p'_{i1}, p'_{i2},... p'_{ij}\}$ where $p_{ij}' = \{m/z_{ij}', I_{ij}'\}$ refers to the $j^{th}$ grouped peak in the $i^{th}$ spectrum.
- The m/z tolerance $t'$ within which two peaks from different spectra are considered to match.

Let the indicator function **I** check whether two peaks from different spectra match within the given tolerance $t'$. For the j$^{th}$ peak in spectrum $i$ and the k$^{th}$ peak in spectrum $l$, we define:

$$I(p'_{ij}, p'_{lk}) = 1 \; if \; |m/z'_{ij} - m/z'_{lk}| \le t'$$
$$= 0 \text{ otherwise}$$

For a given peak, $p'_{ij}$, the number of spectra that contain a matching peak within the tolerance $t'$ is given by:

$$Matches(p'_{ij}) = \sum_{l=1,\, l\neq i}^{N} \sum_{k=1}^{K_l} I(p'_{ij}, p'_{lk})$$ Where $K_l$ is the number of peaks in spectrum $s'_l$.

The **mean m/z** and **mean intensity** for a given peak in the representative spectrum are calculated by taking the average of the m/z values of all matching peaks, including the original peak itself. If the peak $p_{ij}$ has found matches in other spectra, we include those matching peaks in the calculation.

$$Mean(m/z'_{ij}) = \frac{m/z'_{ij} + \sum_{l=1,\, l\neq i}^{N} \sum_{k=1}^{K_l} I(p'_{ij}, p'_{lk}).m/z'_{lk}}{1 + Matches(p'_{ij})}$$

$$Mean(intensity'_{ij}) = \frac{intensity'_{ij} + \sum_{l=1,\, l\neq i}^{N} \sum_{k=1}^{K_l} I(p'_{ij}, p'_{lk}).intensity'_{lk}}{1 + Matches(p'_{ij})}$$

While calculating the aggregate spectra for a given feature, the consistency factor for every representative peak belonging to the aggregate, its mean m/z, and intensity is reported.

The consistency factor $Cf$ for the peak $p'_{ij}$ is the fraction of spectra that contain a matching peak. We can represent $Cf = \frac{1 + Matches(p'_{ij})}{N}$

**Interpretation:**

- The **mean m/z** gives the representative m/z value for a peak across all spectra, adjusted for minor variations within the allowed tolerance.

- The **mean intensity** provides an average intensity across the matching peaks, reflecting the peak's overall abundance across the replicate spectra.

Signal presence count is the number of times a given peak in the representative spectrum is observed across all replicate spectra.

$$Signal\ presence\ count\ =\ Cf\ *\ Number\ of\ spectra$$

**Table S1A: Median values of spectral matching metrics during the validation phase**

| Dataset | Denoising Status | Matching Score (95% CI) | Modified Dot Product (95% CI) | Forward Dot product (95% CI) | Reverse Dot product (95% CI) | Number of Matching fragments (95% CI) | Total experimental fragments (95% CI) | Fragment Matching Ratio (95% CI) |
|---|---|---|---|---|---|---|---|---|
| **WTC-LI (1227 MS2 features)** | **Before** | 580.98 (576.85-589.33) | 0.374 (0.355-0.391) | 0.071 (0.062-0.082) | 0.683 (0.65-0.701) | 17 (16-18) | 314 (302-335.97) | 0.058 (0.056-0.06) |
| | **After** | 673.04 (663.18-678.84) | 0.699 (0.683-0.722) | 0.738 (0.715-0.755) | 0.724 (0.695-0.744) | 11 (10-12) | 21 (19-22) | 0.571 (0.55-0.6) |
| **COVID-19 (941 MS2 features)** | **Before** | 703.31 (694.26-712.35) | 0.581 (0.568-0.595) | 0.592 (0.578-0.606) | 0.597 (0.58-0.615) | 21 (19.2-22.8) | 60 (56.8-63.1) | 0.35 (0.339-0.362) |
| | **After** | 697.21 (684.9-703.23) | 0.649 (0.637-0.661) | 0.672 (0.659-0.684) | 0.666 (0.651-0.681) | 16 (14.5-17.5) | 36 (32.9-37) | 0.471 (0.458-0.483) |
| **ST003050 (526 MS2 features)** | **Before** | 530.542 (521.96-539.12) | 0.663 (0.646-0.679) | 0.666 (0.65-0.683) | 0.744 (0.72-0.769) | 4 (3.69-4.30) | 14 (13.53-14.46) | 0.27 (0.257-0.288) |
| | **After** | 531.659 (524.3-541.2) | 0.726 (0.7-0.736) | 0.775 (0.756-0.789) | 0.739 (0.711-0.75) | 3 (2.8-3.2) | 6 (5.6-6.3) | 0.6 (0.58-0.62) |
| **Mixture of Standards (488 MS2 features)** | **Before** | 620.93 (603.5-638.4) | 0.41 (0.39-0.42) | 0.113 (0.1-0.125) | 0.663 (0.641-0.686) | 18 (13-22) | 197 (142-252) | 0.121 (0.107-0.135) |
| | **After** | 755.052 (736.08 - 774.03) | 0.717 (0.7-0.733) | 0.832 (0.814-0.85) | 0.717 (0.696-0.737) | 14.5 (11.26-17.73) | 29 (24.16-33.83) | 0.69 (0.664-0.715) |

**Table S1B: Number of fragments during three stages of denoising in the validation phase**

| | **Median number of fragments during different denoising stages** | | |
|---|---|---|---|
| **Dataset** | **Before Denoising (95% CI)** | **After intraspectrum grouping (95% CI)** | **After the application of frequency cutoff (95% CI)** |
| **WTC-LI** | 314<br>(302-335.97) | 28<br>(27.3-28.69) | 21<br>(19-21.55) |
| **COVID-19** | 60<br>(56.8-63.1) | 44<br>(42.8-46.9) | 36<br>(32.9-37) |
| **ST003050** | 14<br>(13.53-14.46) | 10<br>(8.9-11.2) | 6<br>(5.6-6.3) |
| **Mixture of Standards** | 197<br>(142-252) | 36<br>(30.15-41.84) | 29<br>(24.16-33.83) |

**Table S2 - Median values of matching metrics across different matching score ranges- (WTC-LI)**

| Before Denoising | | | | | | | |
|---|---|---|---|---|---|---|---|
| **Matching Score Range** | **Number of features** | **Modified Dot Pdt** | **Number of Matching Fragments** | **Total Exp fragments** | **FDP** | **RDP** | **Fragment Matching Ratio** |
| >1000 | 0 | | | | | | |
| 900-1000 | 7 | 0.495 | 126 | 1783 | 0.169 | 0.825 | 0.073 |
| 800-900 | 25 | 0.497 | 58 | 860 | 0.152 | 0.842 | 0.078 |
| 700-800 | 115 | 0.483 | 35 | 568 | 0.089 | 0.804 | 0.069 |
| 600-700 | 333 | 0.375 | 23 | 437 | 0.045 | 0.686 | 0.064 |
| 500-600 | 439 | 0.328 | 15 | 293 | 0.024 | 0.592 | 0.057 |
| <500 | 229 | 0.394 | 6 | 133 | 0.013 | 0.769 | 0.043 |
| **After Denoising** | | | | | | | |
| >1000 | 17 | 0.939 | 81 | 92 | 0.948 | 0.926 | 0.883 |
| 900-1000 | 49 | 0.909 | 31 | 40 | 0.913 | 0.895 | 0.8 |
| 800-900 | 148 | 0.881 | 18 | 26 | 0.898 | 0.871 | 0.74 |
| 700-800 | 269 | 0.727 | 14 | 26 | 0.75 | 0.771 | 0.633 |
| 600-700 | 434 | 0.629 | 10 | 20 | 0.678 | 0.636 | 0.527 |
| 500-600 | 240 | 0.716 | 4 | 11 | 0.743 | 0.71 | 0.449 |
| <500 | 70 | 0.533 | 4 | 10 | 0.461 | 0.516 | 0.333 |

**Table S3 - Median values of matching metrics across different matching score ranges- (COVID-19)**

| Before Denoising | | | | | | | |
|---|---|---|---|---|---|---|---|
| **Matching Score Range** | **Number of features** | **Modified Dot Pdt** | **Number of Matching Fragments** | **Total Exp fragments** | **FDP** | **RDP** | **Fragment Matching Ratio** |
| >1000 | 34 | 0.814 | 100.5 | 179 | 0.781 | 0.875 | 0.596 |
| 900-1000 | 62 | 0.807 | 38.5 | 103.5 | 0.756 | 0.854 | 0.46 |
| 800-900 | 111 | 0.72 | 31 | 80 | 0.669 | 0.782 | 0.425 |
| 700-800 | 251 | 0.517 | 28 | 69 | 0.523 | 0.536 | 0.375 |
| 600-700 | 275 | 0.511 | 15 | 47 | 0.593 | 0.481 | 0.304 |
| 500-600 | 131 | 0.557 | 7 | 41 | 0.482 | 0.571 | 0.194 |
| <500 | 31 | 0.493 | 4 | 30 | 0.41 | 0.559 | 0.133 |
| **After Denoising** | | | | | | | |
| >1000 | 35 | 0.866 | 91 | 117 | 0.873 | 0.903 | 0.799 |
| 900-1000 | 58 | 0.832 | 35 | 55 | 0.823 | 0.869 | 0.676 |
| 800-900 | 118 | 0.748 | 28 | 48 | 0.723 | 0.784 | 0.593 |
| 700-800 | 247 | 0.579 | 24 | 47 | 0.587 | 0.587 | 0.51 |
| 600-700 | 272 | 0.6 | 11 | 27 | 0.686 | 0.561 | 0.429 |
| 500-600 | 168 | 0.631 | 5 | 21 | 0.659 | 0.602 | 0.295 |
| <500 | 43 | 0.599 | 3 | 15 | 0.468 | 0.646 | 0.25 |

**Table S4 - Median values of matching metrics across different matching score ranges- (ST003050)**

| Before Denoising | | | | | | | |
|---|---|---|---|---|---|---|---|
| Matching Score Range | Number of features | Modified Dot Pdt | Number of Matching Fragments | Total Exp fragments | FDP | RDP | Fragment Matching Ratio |
| >1000 | | | | | | | |
| 900-1000 | | | | | | | |
| 800-900 | 1 | 0.91 | 14 | 26 | 0.882 | 0.938 | 0.538 |
| 700-800 | 10 | 0.893 | 7 | 17 | 0.813 | 0.977 | 0.442 |
| 600-700 | 134 | 0.836 | 5 | 14 | 0.808 | 0.906 | 0.425 |
| 500-600 | 187 | 0.725 | 3 | 15 | 0.734 | 0.818 | 0.267 |
| <500 | 194 | 0.541 | 3 | 14 | 0.488 | 0.544 | 0.2 |
| **After Denoising** | | | | | | | |
| >1000 | | | | | | | |
| 900-1000 | | | | | | | |
| 800-900 | 2 | 0.944 | 11.5 | 13.5 | 0.925 | 0.963 | 0.85 |
| 700-800 | 11 | 0.922 | 7 | 9 | 0.922 | 0.959 | 0.889 |
| 600-700 | 133 | 0.922 | 4 | 5 | 0.923 | 0.924 | 0.8 |
| 500-600 | 168 | 0.81 | 3 | 5 | 0.832 | 0.807 | 0.6 |
| <500 | 232 | 0.607 | 2 | 6 | 0.63 | 0.552 | 0.4 |

**Table S5 - Median values of matching metrics across different matching score ranges-(benchmarking)**

| Before Denoising | | | | | | | |
|---|---|---|---|---|---|---|---|
| **Matching Score Range** | **Number of features** | **Modified Dot Pdt** | **Number of Matching Fragments** | **Total Exp fragments** | **FDP** | **RDP** | **Fragment Matching Ratio** |
| >1000 | 1 | 0.448 | 235 | 2045 | 0.105 | 0.792 | 0.114 |
| 900-1000 | 17 | 0.449 | 136 | 1310 | 0.101 | 0.719 | 0.117 |
| 800-900 | 56 | 0.421 | 97 | 751 | 0.101 | 0.71 | 0.126 |
| 700-800 | 76 | 0.457 | 38 | 311 | 0.128 | 0.755 | 0.136 |
| 600-700 | 83 | 0.463 | 21 | 226 | 0.122 | 0.769 | 0.118 |
| 500-600 | 76 | 0.405 | 12 | 110 | 0.108 | 0.673 | 0.127 |
| <500 | 137 | 0.374 | 4 | 45 | 0.105 | 0.557 | 0.095 |
| **After Denoising** | | | | | | | |
| >1000 | 49 | 0.846 | 97 | 127 | 0.89 | 0.843 | 0.801 |
| 900-1000 | 76 | 0.86 | 40 | 60.5 | 0.864 | 0.846 | 0.738 |
| 800-900 | 88 | 0.856 | 20 | 33 | 0.868 | 0.845 | 0.746 |
| 700-800 | 70 | 0.702 | 15 | 25 | 0.799 | 0.695 | 0.696 |
| 600-700 | 54 | 0.693 | 8.5 | 16.5 | 0.754 | 0.697 | 0.667 |
| 500-600 | 55 | 0.703 | 4 | 10 | 0.72 | 0.712 | 0.667 |
| <500 | 96 | 0.568 | 2 | 6.5 | 0.563 | 0.47 | 0.348 |

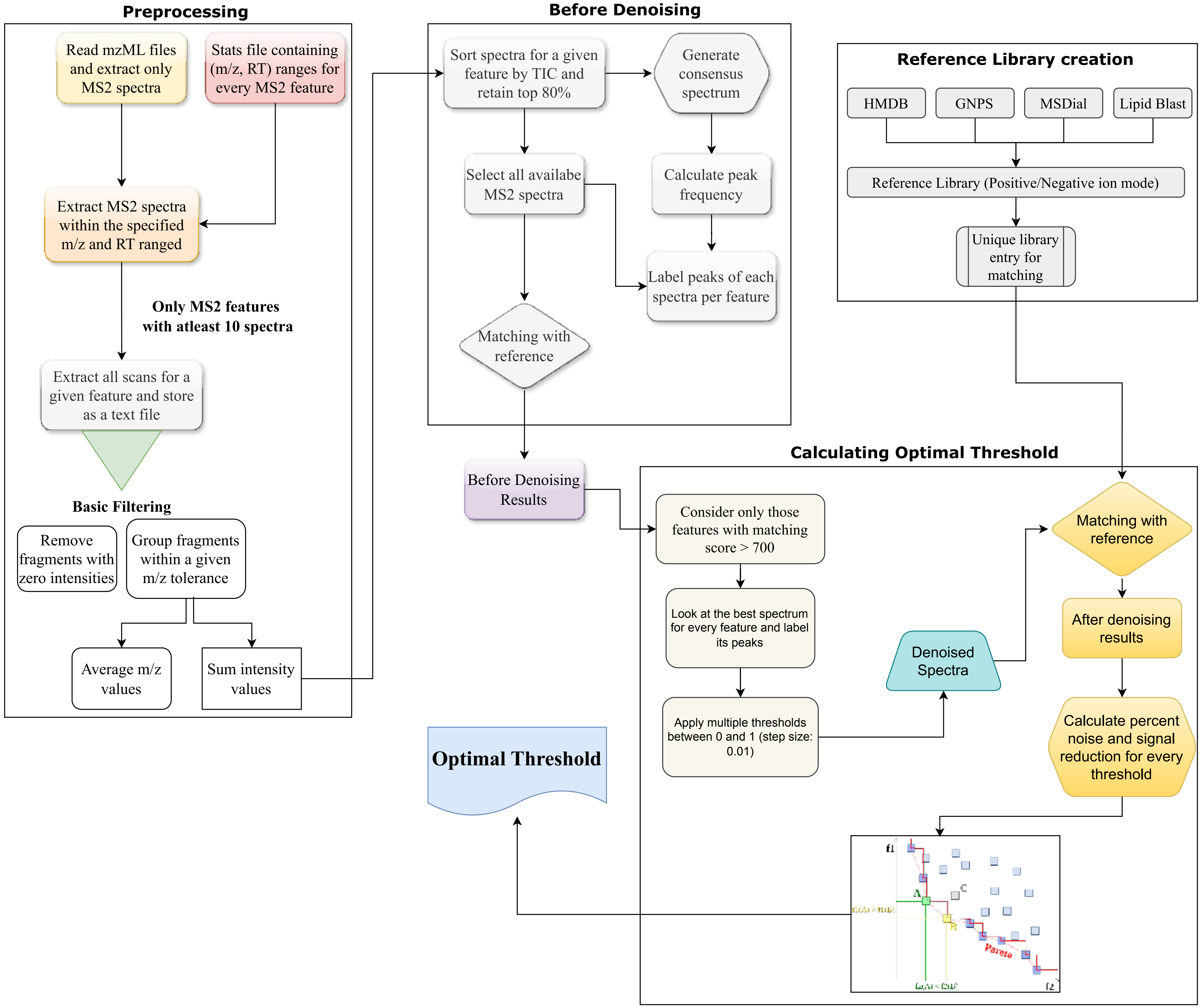


**Figure S1: Tuning phase workflow to derive optimal frequency threshold**

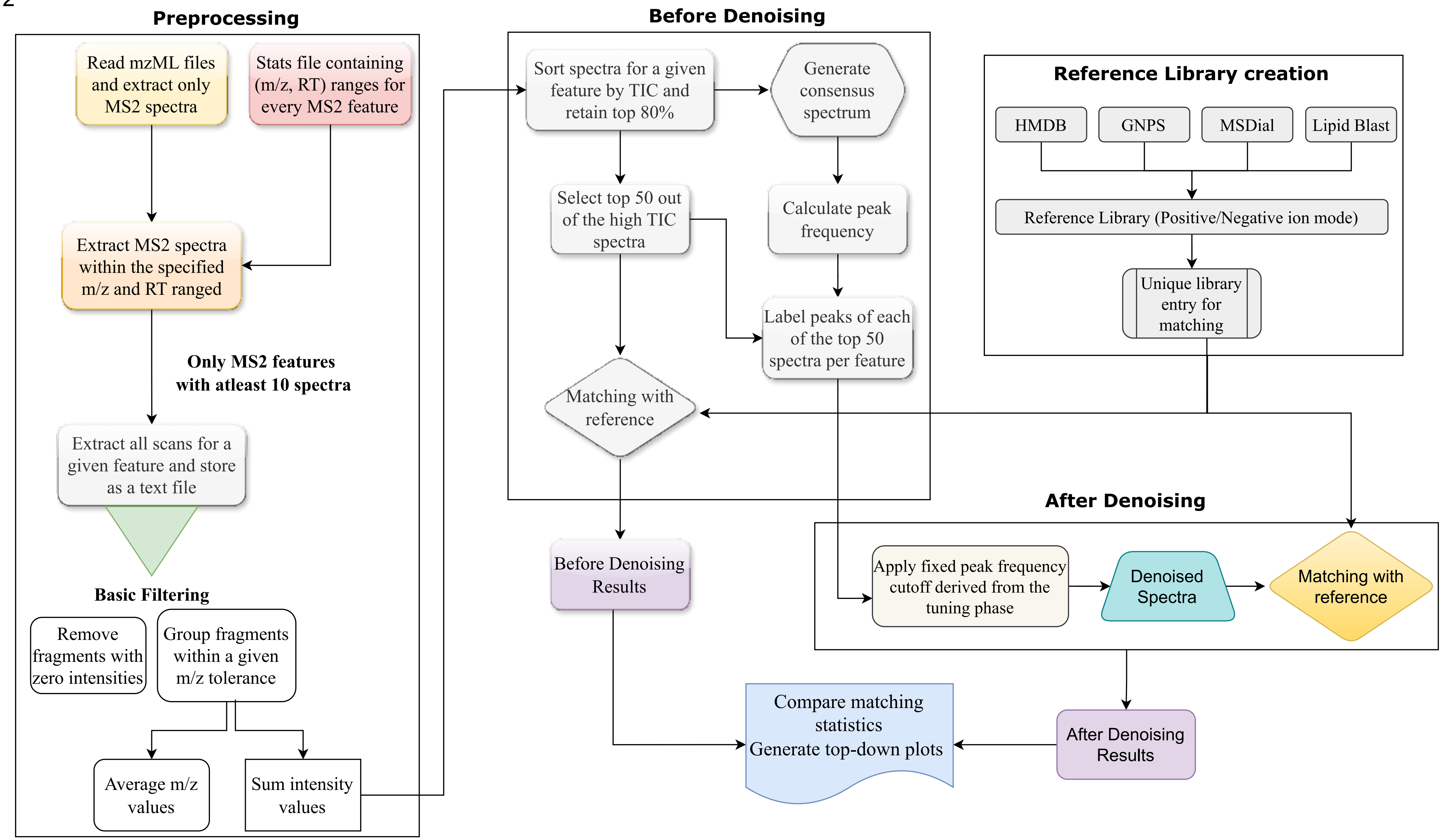


**Figure S2: Testing phase workflow to evaluate the optimal frequency threshold**

Forward Dot Product =

$$\frac{20*40 + 30*20 + 40*45 + 5*0 + 25*25 + 2*0 + 5*0 + 20*15 + 30*40 + 6*0 + 60*45}{|Experimental\_fwd| * |Library\_fwd|}$$

Experimental_fwd = (20, 30, 40, 5, 25, 2, 5, 20, 30, 6, 60)      Library_fwd = (40, 20, 45, 0, 25, 0, 0, 15, 40, 0, 45)

**Modified Dot Product = 0.5 * [ Forward Dot Product + Reverse Dot Product]**

**Matching Score = 5 * [100* Modified Dot Product + 20 * $\log_2$(Number of Matching Fragments)]**

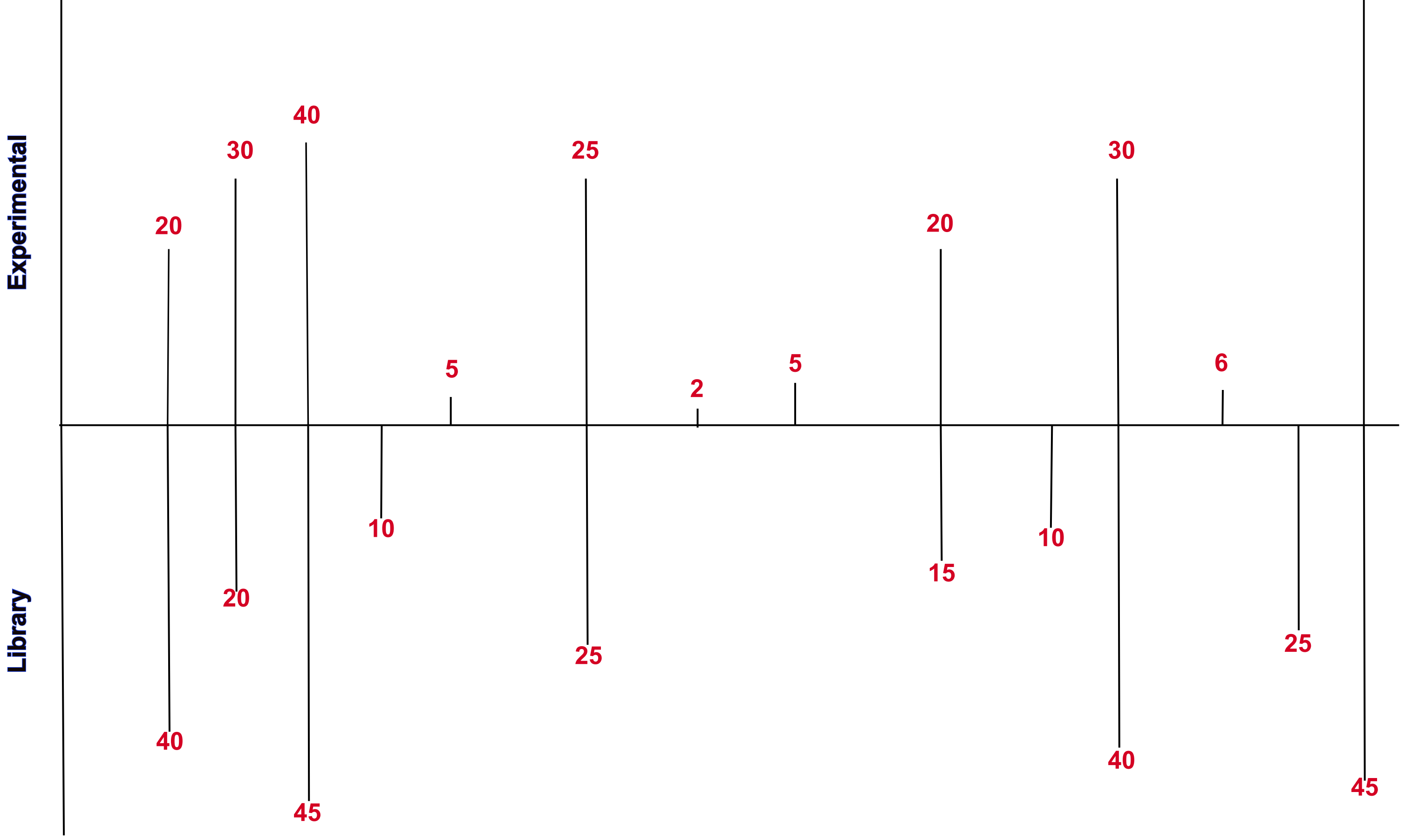


Reverse Dot Product =

$$\frac{20*40 + 30*20 + 40*45 + 0*10 + 5*0 +25*25 + 2*0 + 5*0 + 20*15 + 0.*10 + 30*40 + 6*0 + 0* 25 + 60*45}{|Experimental\_rvr| * |Library\_rvr|}$$

Experimental_rvr = (20, 30, 40, 0, 25, 20, 0, 30, 0, 60)      Library_rvr = (40, 20, 45, 10, 25, 15, 10, 40, 25, 45)

**Figure S3 - Graphical representation and formulation of matching metrics used in this study**

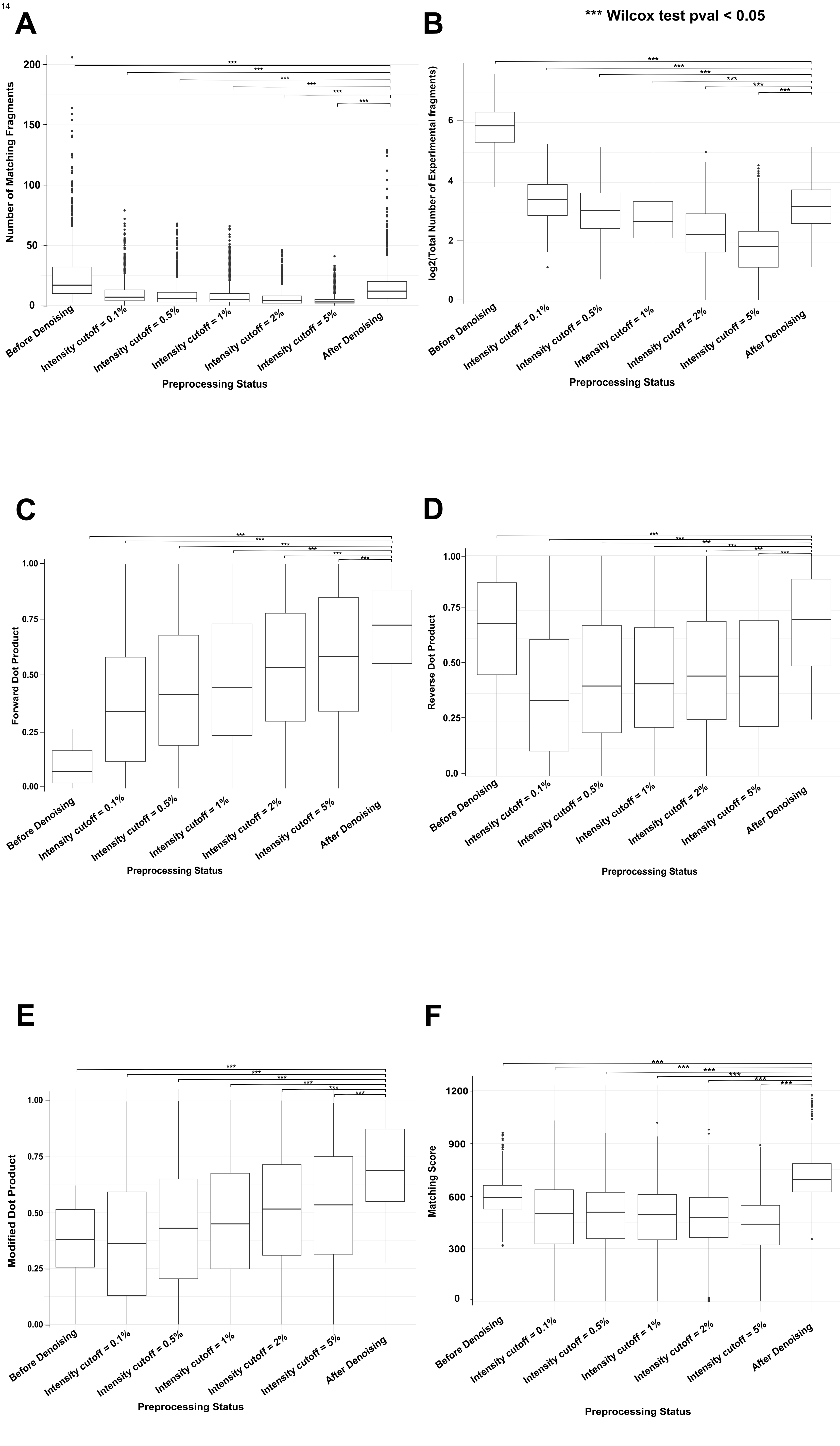


**Figure S4 (i): Comparison of matching metrics between different denoising approaches using the WTC-LI dataset**

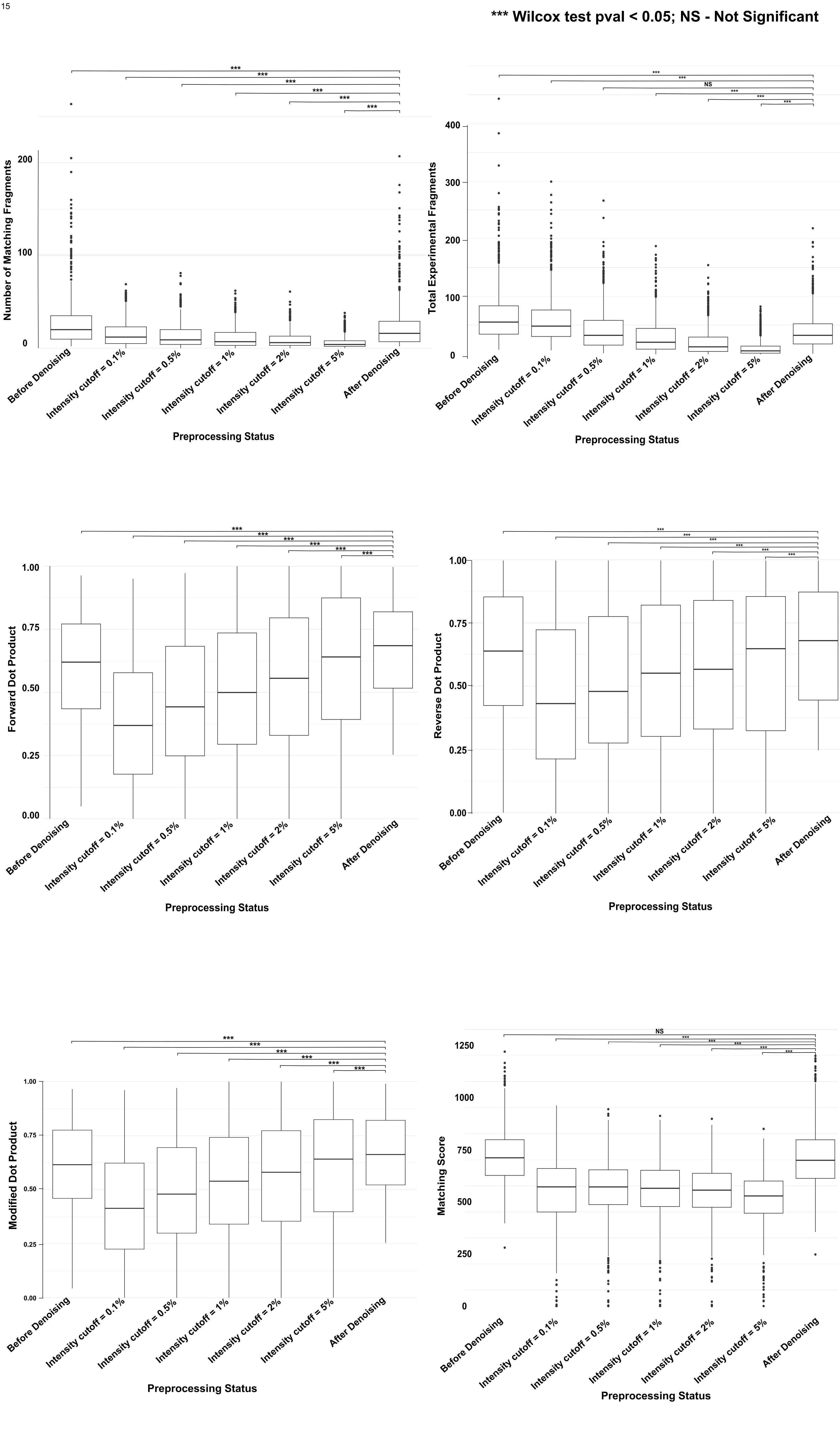


**Figure S4 (ii): Comparison of matching metrics between different denoising approaches using the COVID-19 dataset**

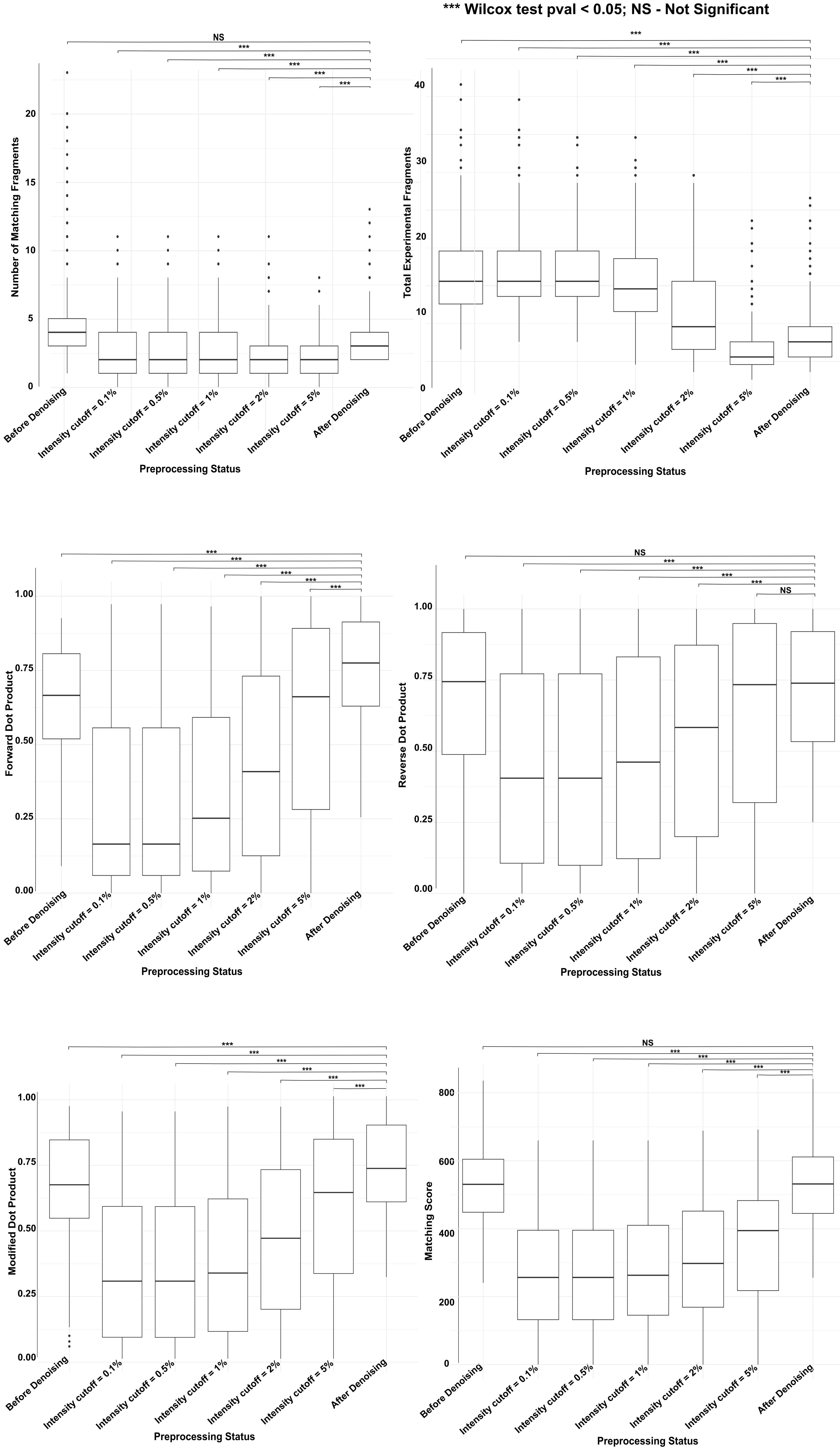


**Figure S4 (iii): Comparison of matching metrics between different denoising approaches using the ST003050 dataset**

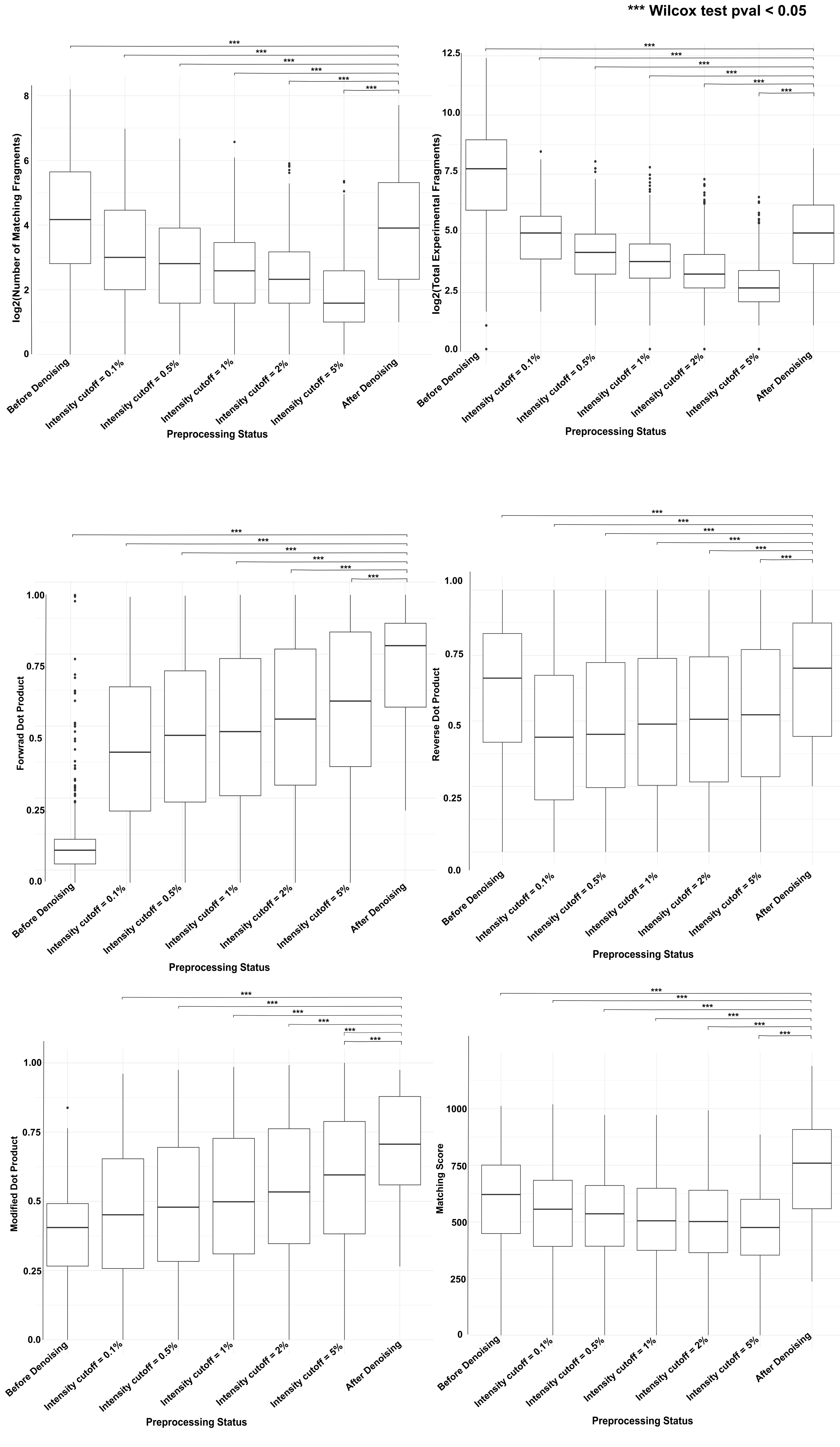


**Figure S4 (iv): Comparison of matching metrics between different denoising approaches using the benchmarking dataset**

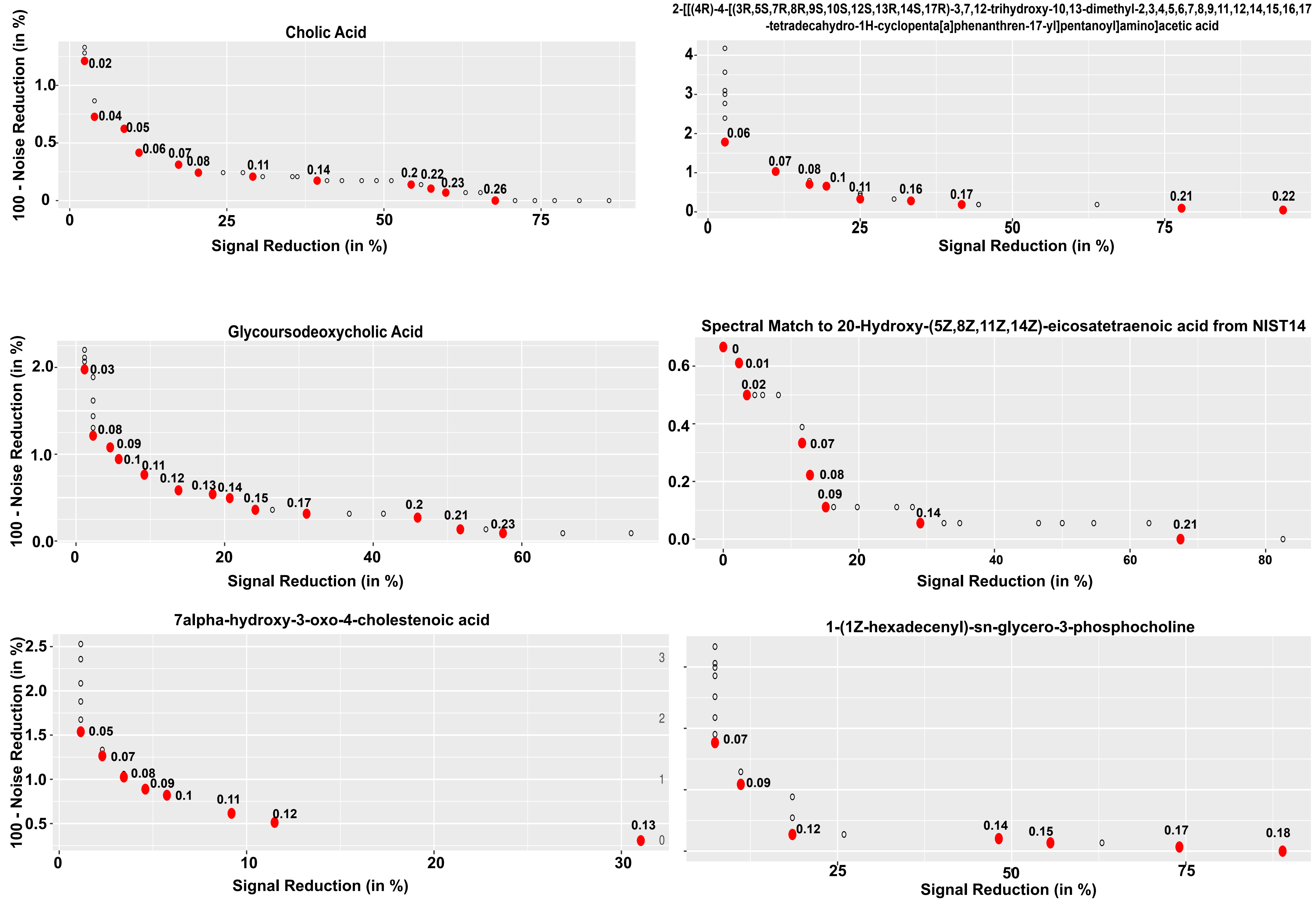


**Figure S5: Plots showing the tradeoff between signal and noise reduction in the tuning phase. The optimal frequencies on the pareto front are shown in red. The maximum matching score among all the Pareto optimal solutions was chosen as the single optimal solution.**

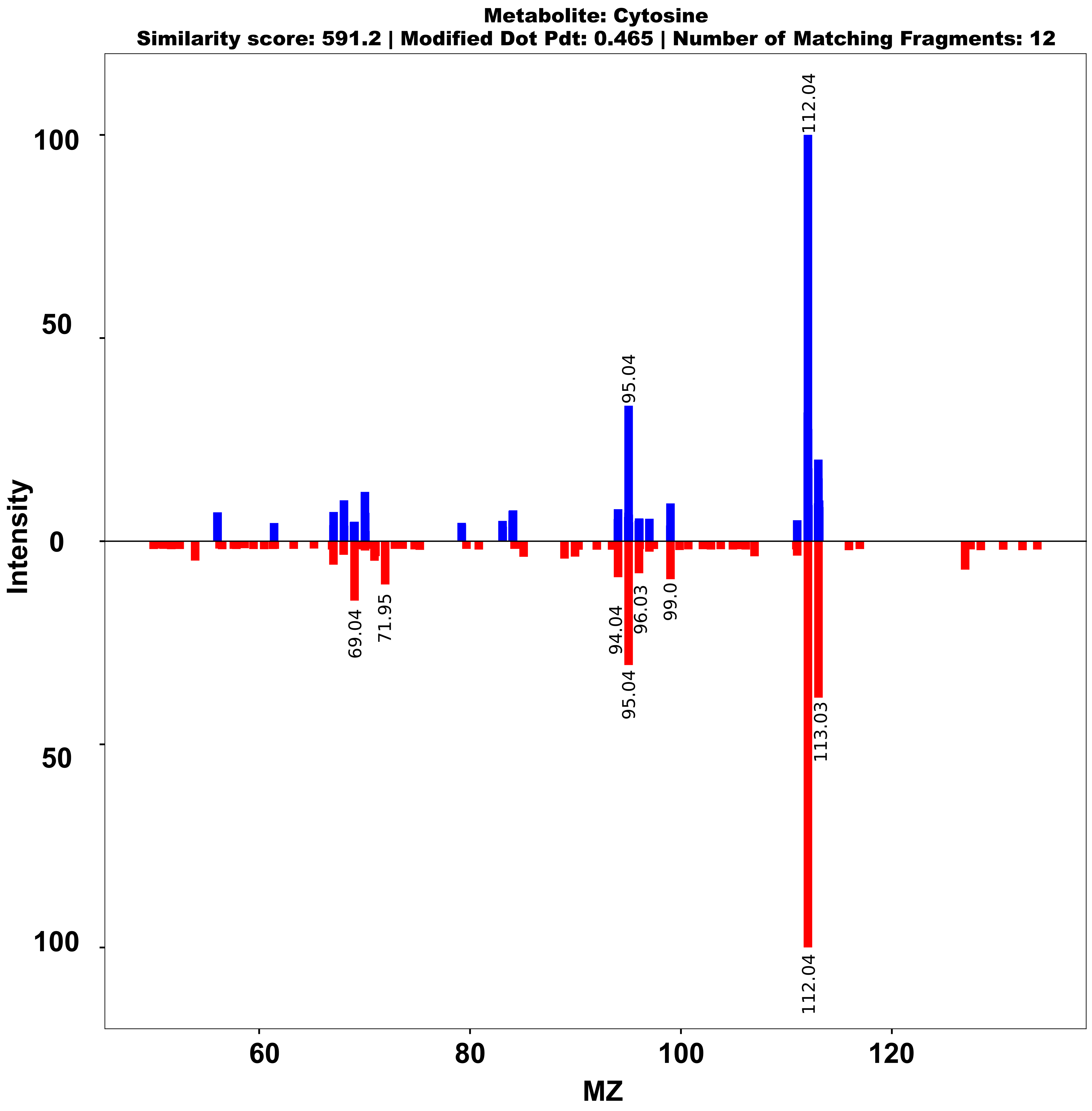
Metabolite: Cytosine
Similarity score: 591.2 | Modified Dot Pdt: 0.465 | Number of Matching Fragments: 12
Intensity
MZ
112.04
95.04
69.04
71.95
94.04
96.03
99.0
95.04
113.03
112.04

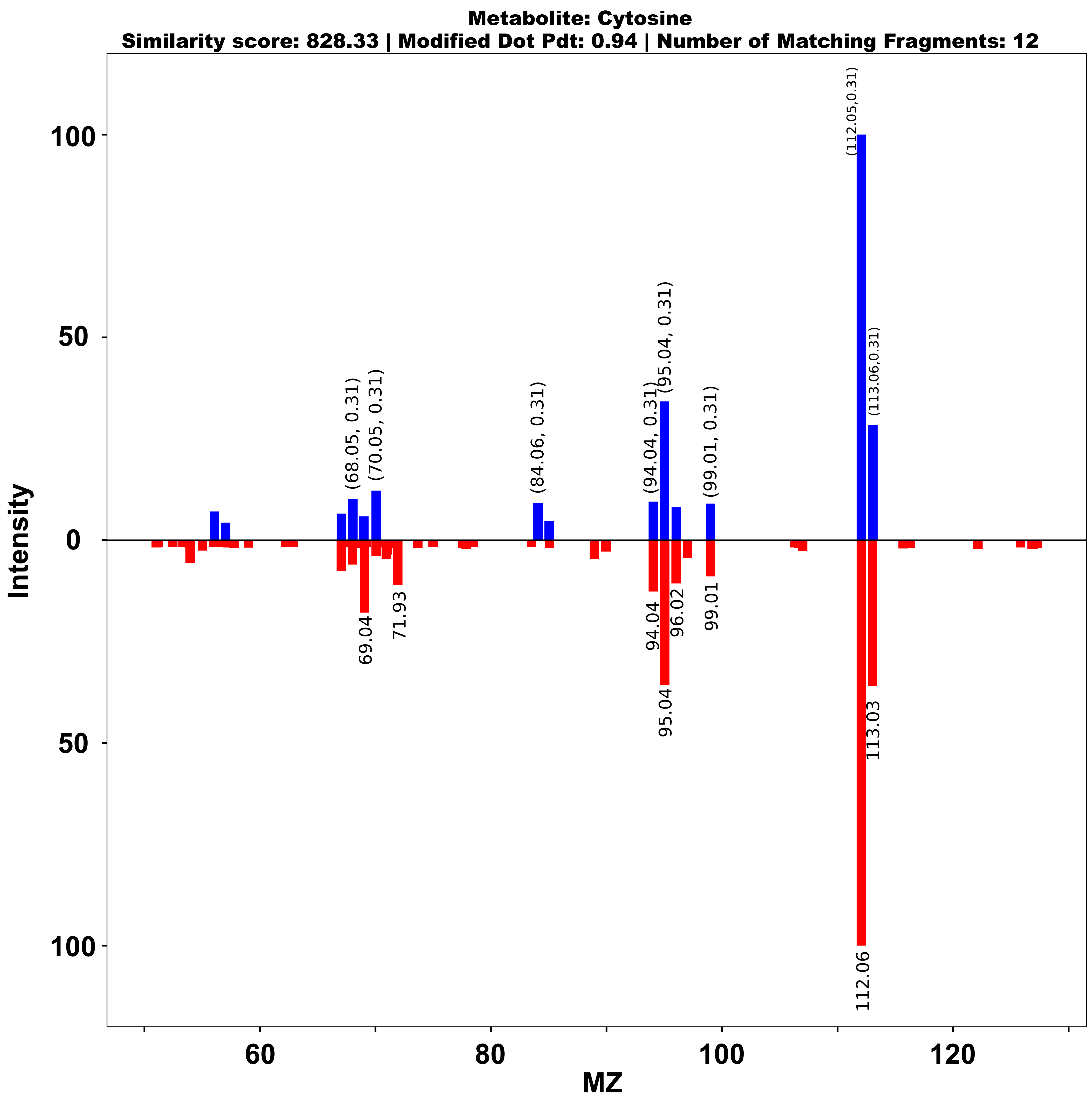
Metabolite: Cytosine
Similarity score: 828.33 | Modified Dot Pdt: 0.94 | Number of Matching Fragments: 12
Intensity
MZ
(112.05,0.31)
(113.06,0.31)
(68.05, 0.31)
(70.05, 0.31)
(84.06, 0.31)
(94.04, 0.31)
(95.04, 0.31)
(99.01, 0.31)
69.04
71.93
94.04
96.02
99.01
95.04
113.03
112.06

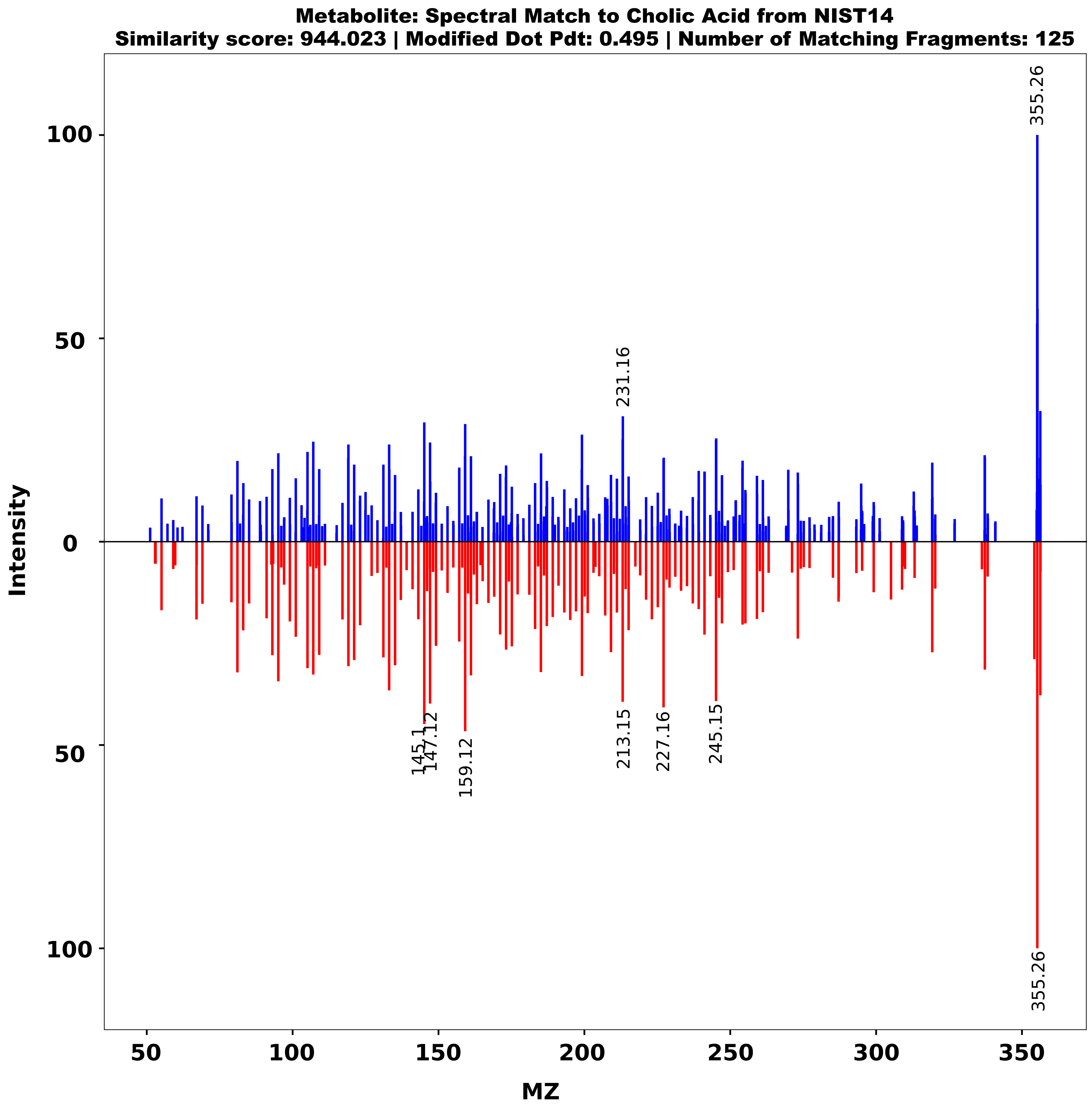

Metabolite: Spectral Match to Cholic Acid from NIST14
Similarity score: 944.023 | Modified Dot Pdt: 0.495 | Number of Matching Fragments: 125
Intensity
MZ
355.26
231.16
145.1
147.12
159.12
213.15
227.16
245.15
355.26


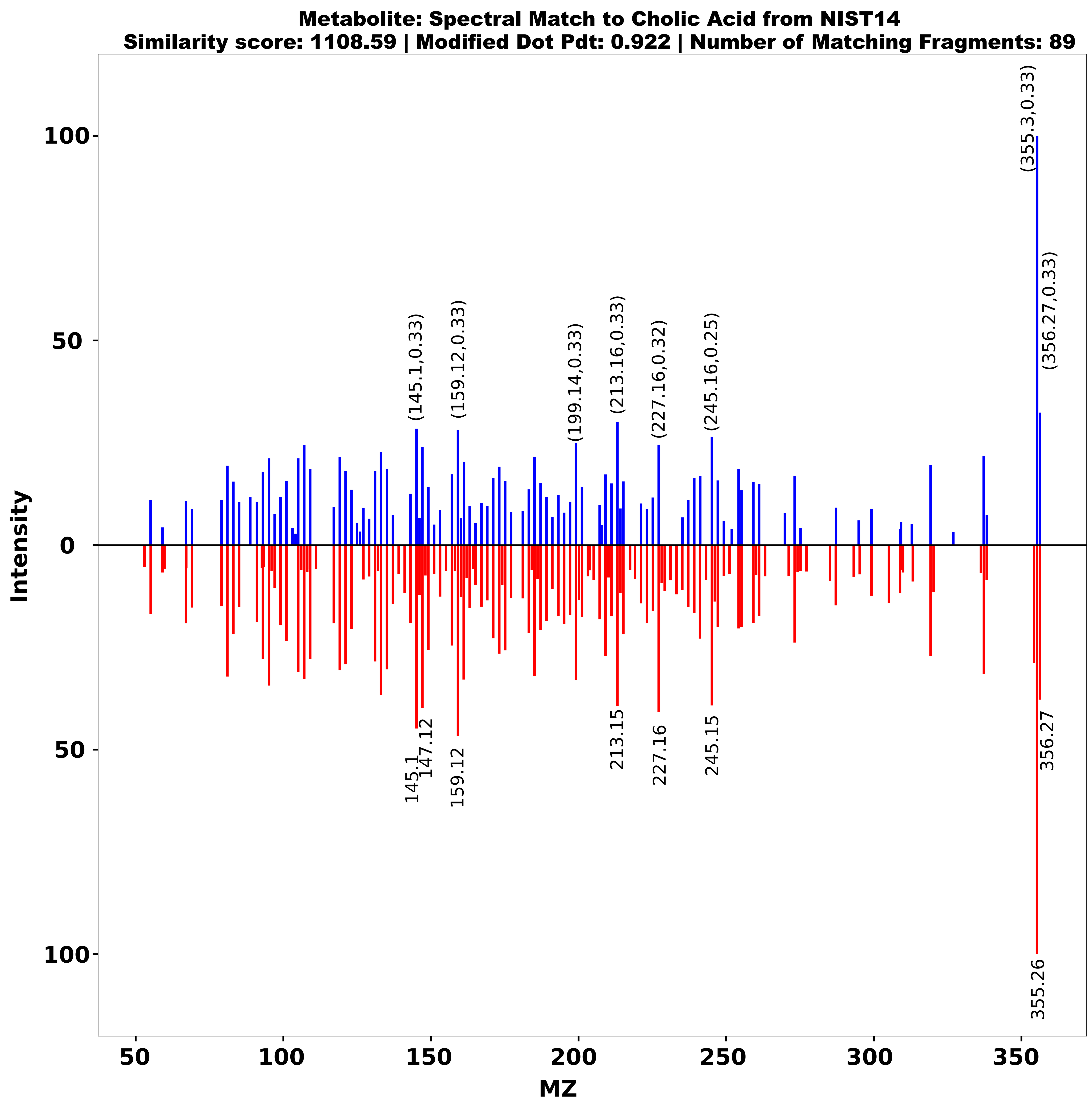

Metabolite: Spectral Match to Cholic Acid from NIST14
Similarity score: 1108.59 | Modified Dot Pdt: 0.922 | Number of Matching Fragments: 89
Intensity
MZ
(355.3,0.33)
(356.27,0.33)
(145.1,0.33)
(159.12,0.33)
(199.14,0.33)
(213.16,0.33)
(227.16,0.32)
(245.16,0.25)
145.1
147.12
159.12
213.15
227.16
245.15
356.27
355.26

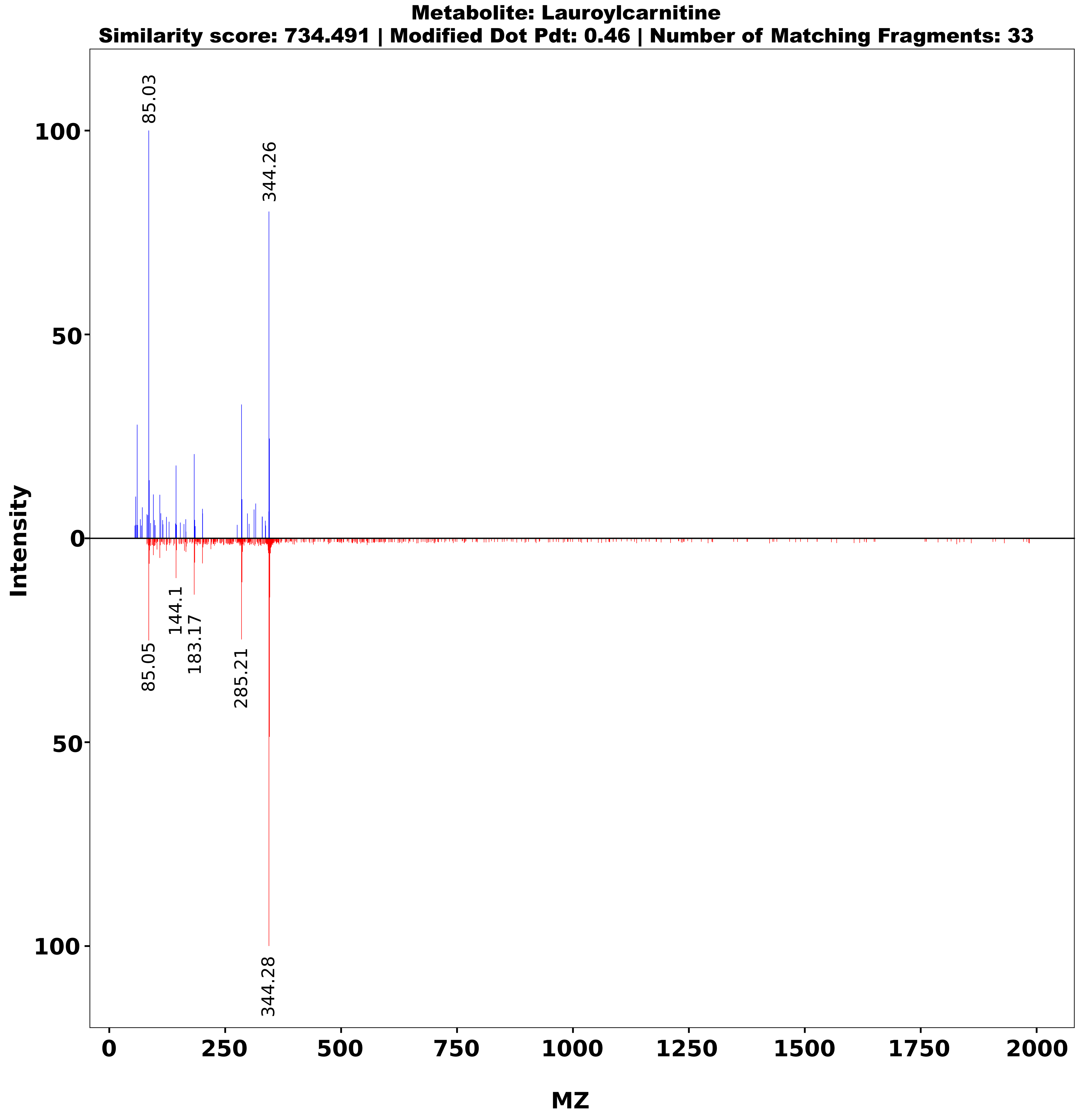
Metabolite: Lauroylcarnitine
Similarity score: 734.491 | Modified Dot Pdt: 0.46 | Number of Matching Fragments: 33
Intensity
MZ
85.03
344.26
85.05
144.1
183.17
285.21
344.28

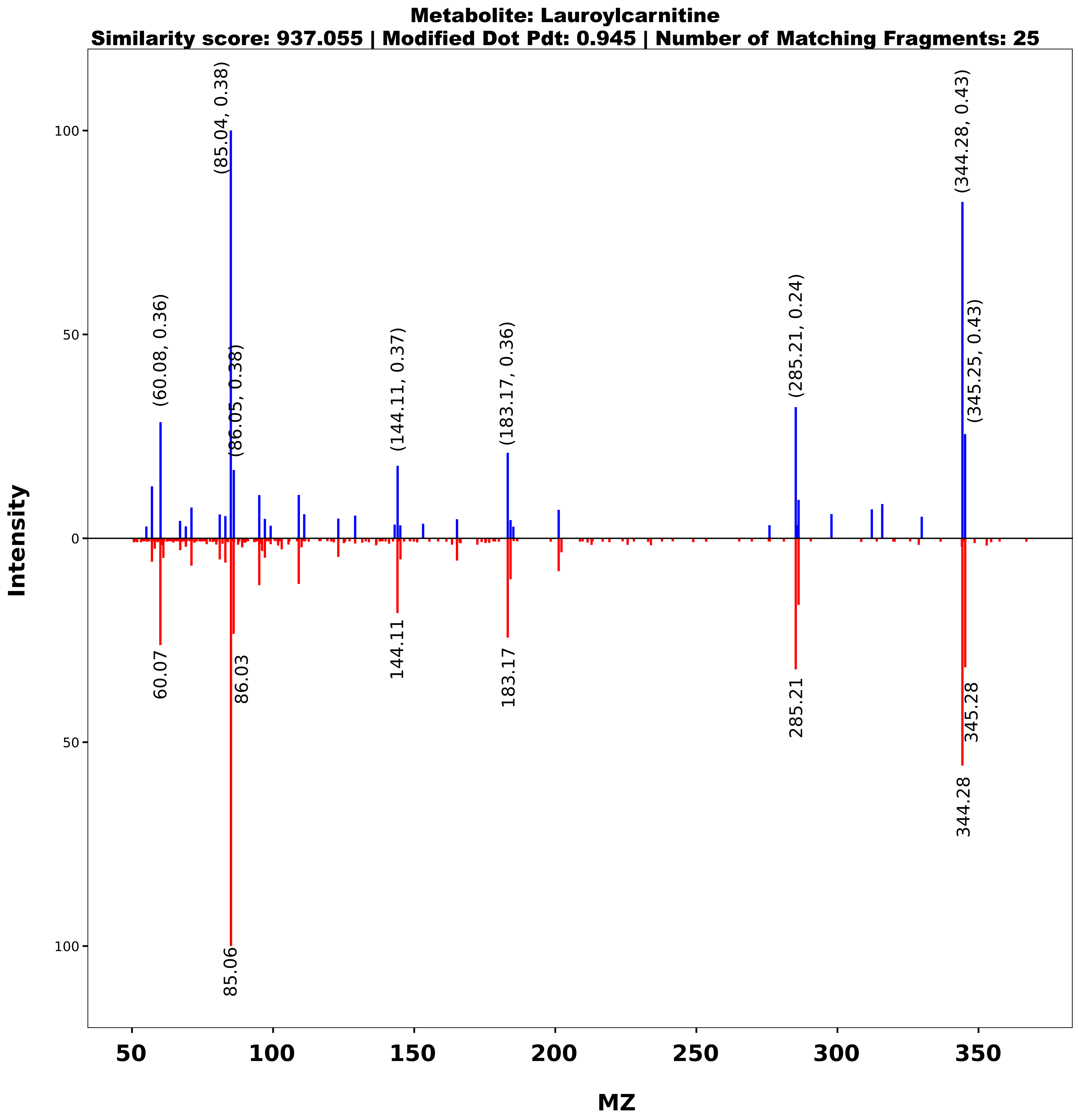
Metabolite: Lauroylcarnitine
Similarity score: 937.055 | Modified Dot Pdt: 0.945 | Number of Matching Fragments: 25
Intensity
MZ
(85.04, 0.38)
(60.08, 0.36)
(86.05, 0.38)
(144.11, 0.37)
(183.17, 0.36)
(285.21, 0.24)
(344.28, 0.43)
(345.25, 0.43)
60.07
86.03
144.11
183.17
285.21
345.28
344.28
85.06

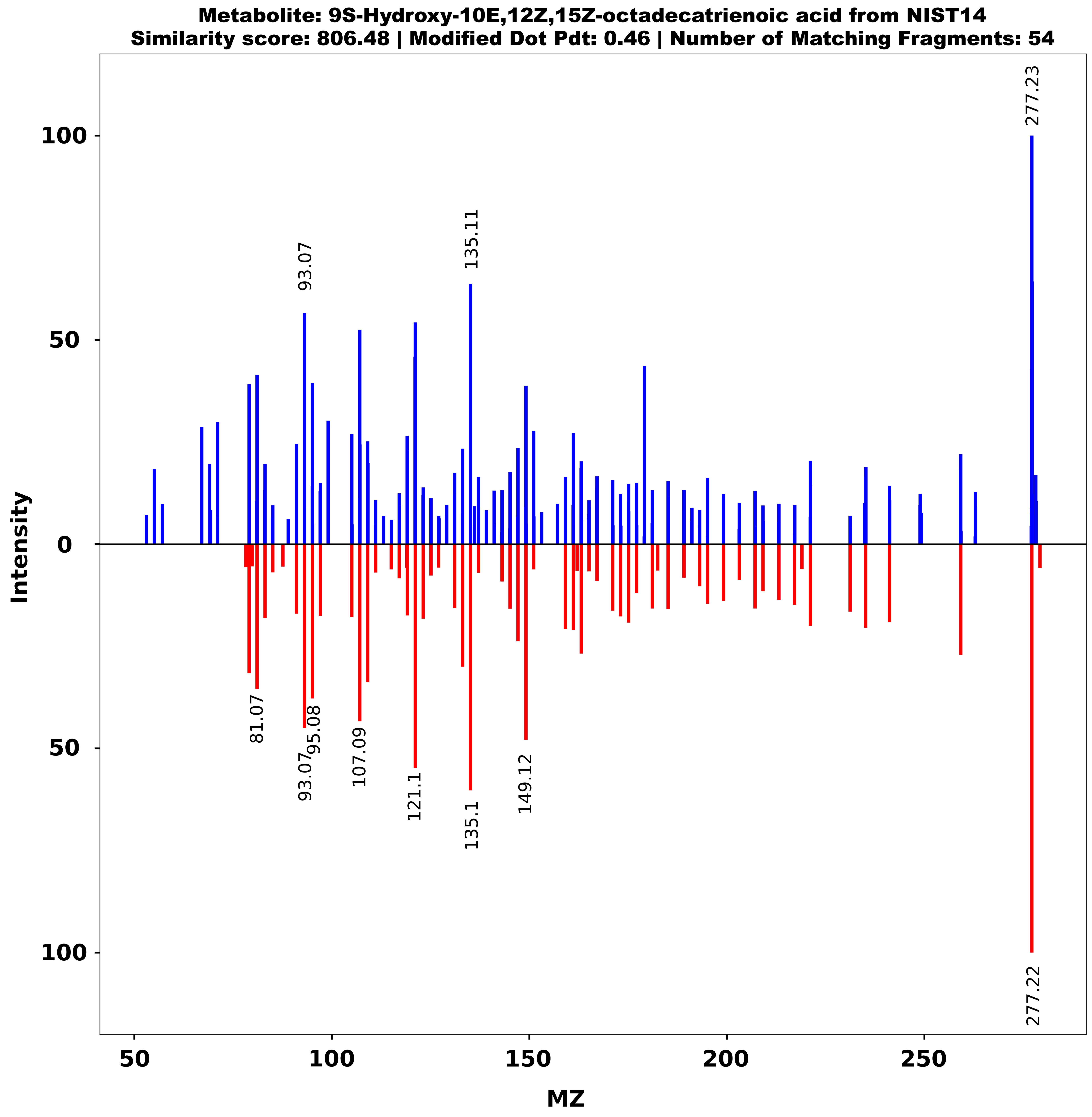
Metabolite: 9S-Hydroxy-10E,12Z,15Z-octadecatrienoic acid from NIST14
Similarity score: 806.48 | Modified Dot Pdt: 0.46 | Number of Matching Fragments: 54
Intensity
MZ
277.23
93.07
135.11
81.07
93.07
95.08
107.09
121.1
135.1
149.12
277.22

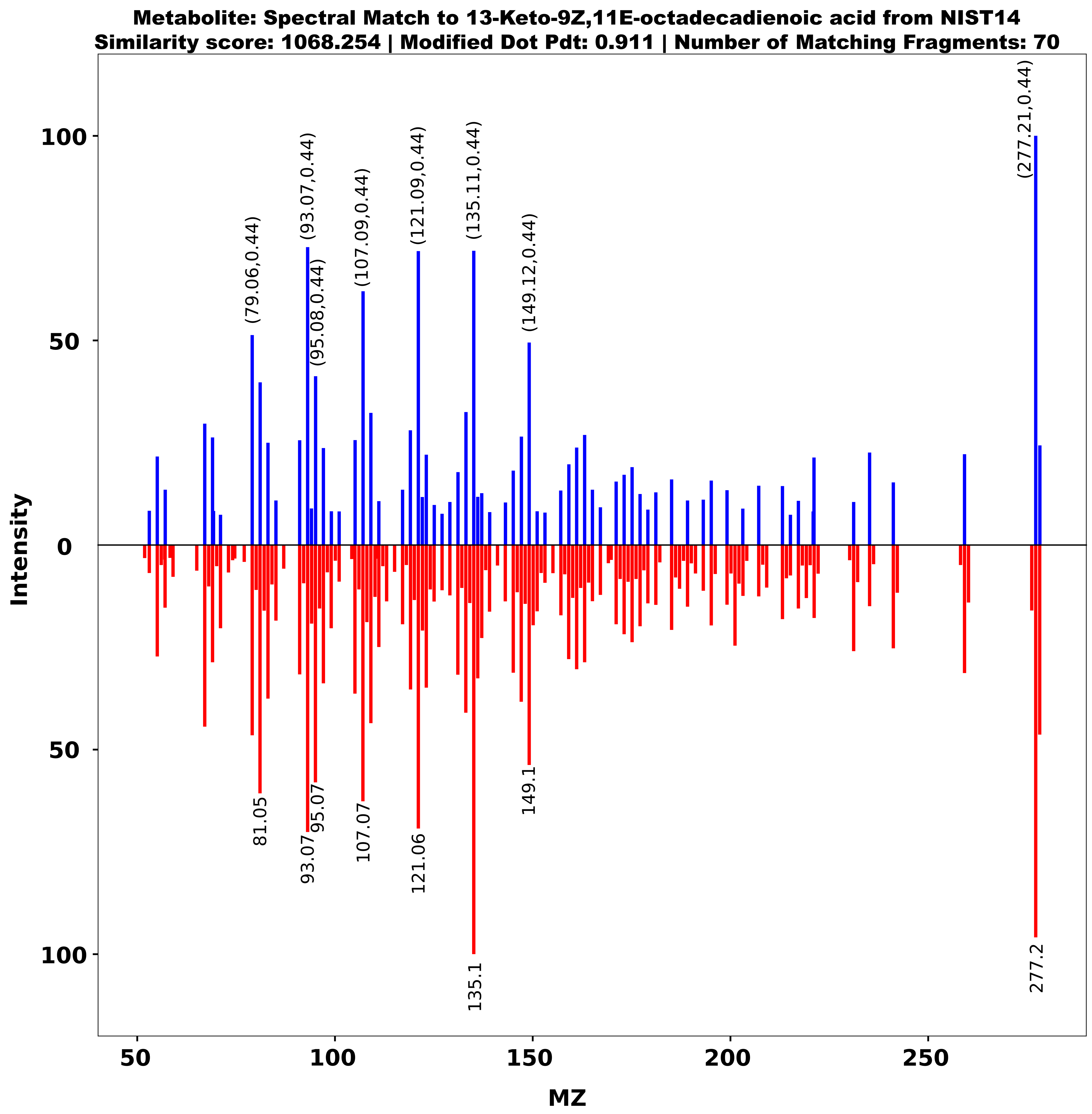
Metabolite: Spectral Match to 13-Keto-9Z,11E-octadecadienoic acid from NIST14
Similarity score: 1068.254 | Modified Dot Pdt: 0.911 | Number of Matching Fragments: 70
Intensity
MZ
(79.06,0.44)
(93.07,0.44)
(95.08,0.44)
(107.09,0.44)
(121.09,0.44)
(135.11,0.44)
(149.12,0.44)
(277.21,0.44)
81.05
93.07
95.07
107.07
121.06
135.1
149.1
277.2

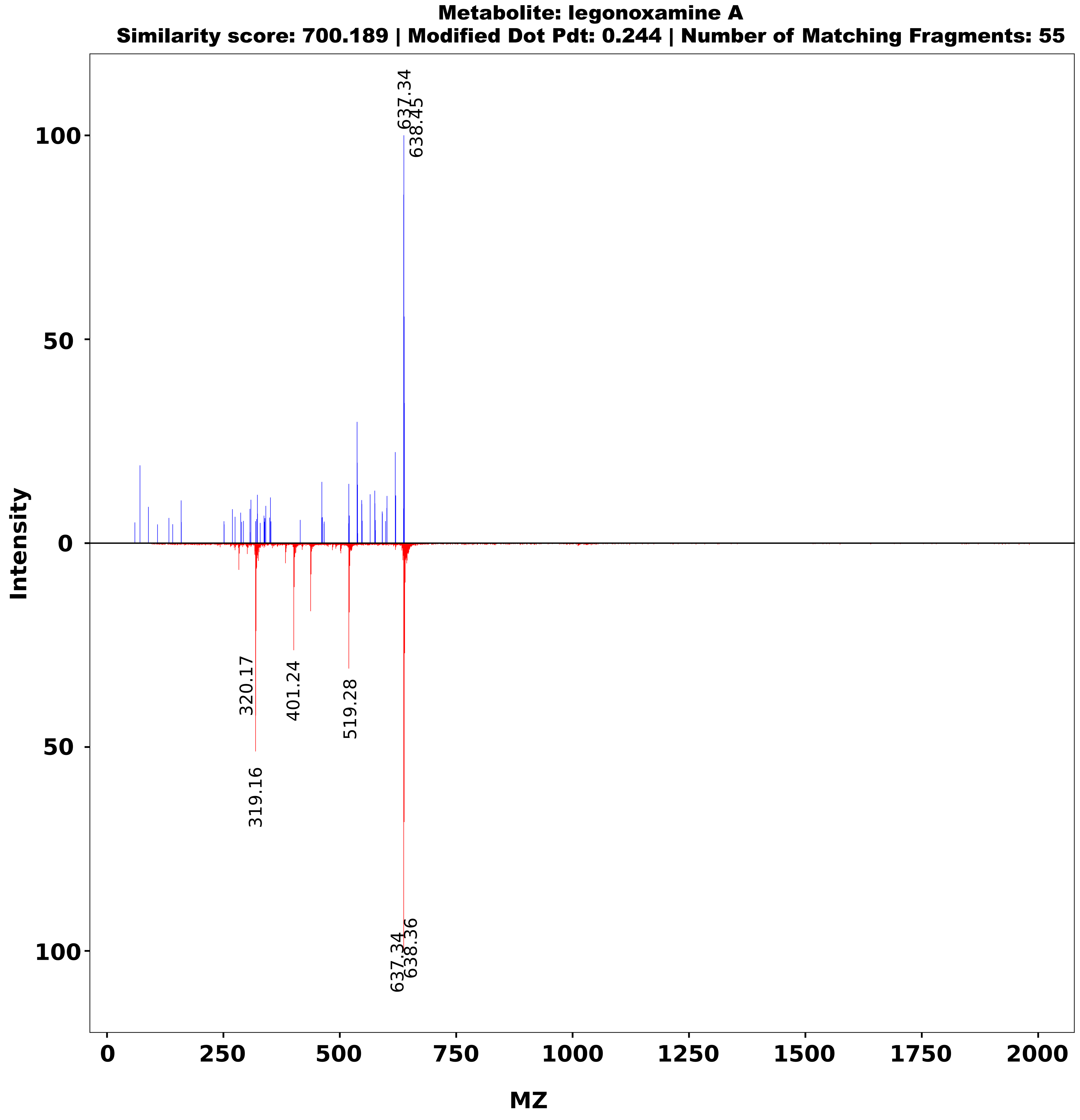
Metabolite: legonoxamine A
Similarity score: 700.189 | Modified Dot Pdt: 0.244 | Number of Matching Fragments: 55
Intensity
MZ
637.34
638.45
320.17
401.24
519.28
319.16
637.34
638.36

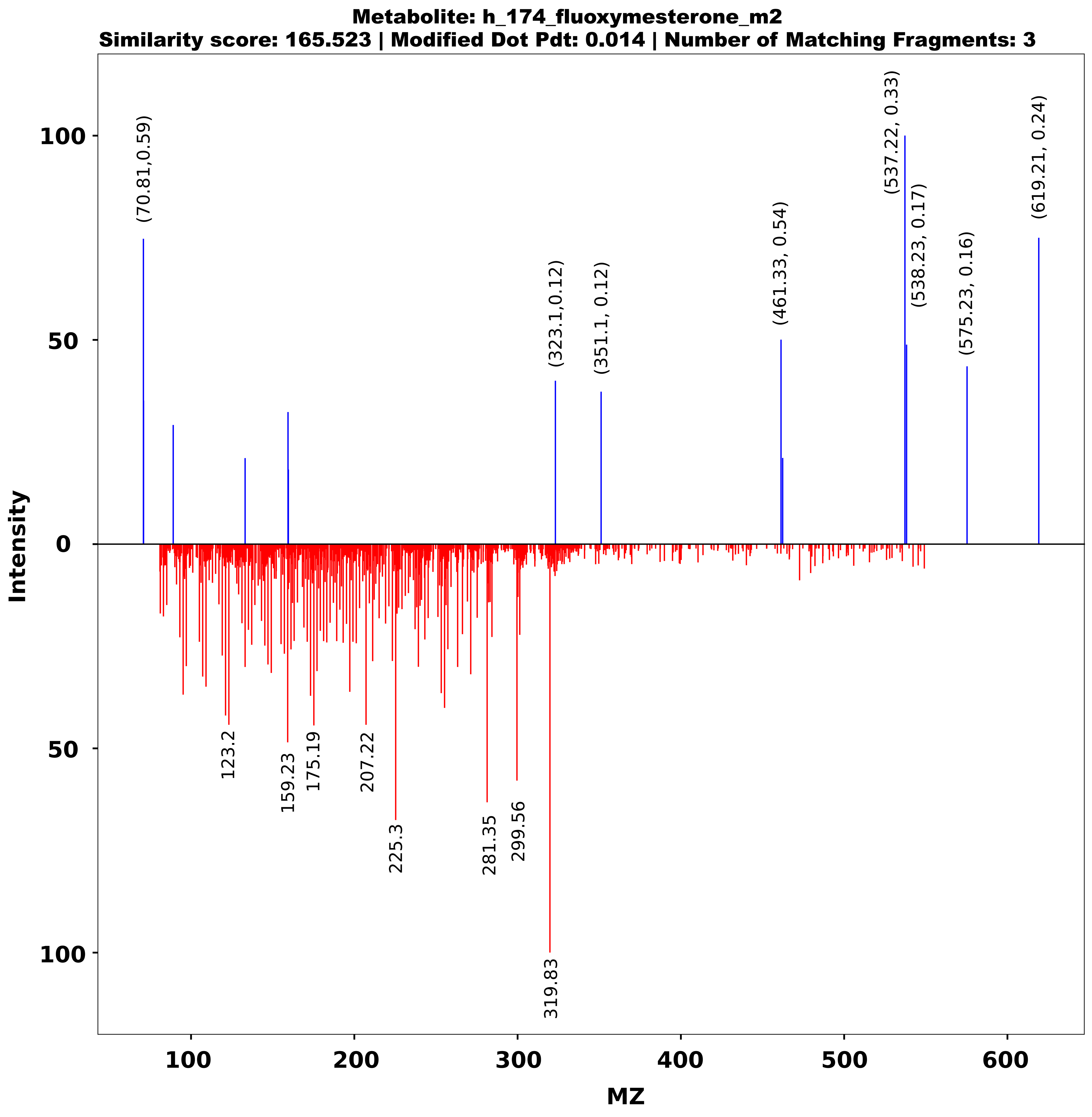
Metabolite: h_174_fluoxymesterone_m2
Similarity score: 165.523 | Modified Dot Pdt: 0.014 | Number of Matching Fragments: 3
Intensity
MZ
(70.81,0.59)
(323.1,0.12)
(351.1, 0.12)
(461.33, 0.54)
(537.22, 0.33)
(538.23, 0.17)
(575.23, 0.16)
(619.21, 0.24)
123.2
159.23
175.19
207.22
225.3
281.35
299.56
319.83
